# Post-EMT: Cadherin-11 mediates cancer hijacking fibroblasts

**DOI:** 10.1101/729491

**Authors:** Weirong Kang, Yibo Fan, Yinxiao Du, Elina A. Tonkova, Yi-Hsin Hsu, Kel Vin Tan, Stephanie Alexander, Bin Sheng Wong, Haocheng Yang, Jingyuan Luo, Kuo Yao, Jiayao Yang, Xin Hu, Tingting Liu, Yu Gan, Jian Zhang, Jean J. Zhao, Konstantinos Konstantopoulos, Peter Friedl, Pek Lan Khong, Aiping Lu, Mien-Chie Hung, Michael B. Brenner, Jeffrey E. Segall, Zhizhan Gu

## Abstract

Current prevailing knowledge on EMT (epithelial mesenchymal transition) deems epithelial cells acquire the characters of mesenchymal cells to be capable of invading and metastasizing on their own. One of the signature events of EMT is called “cadherin switch”, e.g. the epithelial E-cadherin switching to the mesenchymal Cadherin-11. Here, we report the critical events after EMT that cancer cells utilize cadherin-11 to hijack the endogenous cadherin-11 positive fibroblasts. Numerous 3-D cell invasion assays with high-content live cell imaging methods reveal that cadherin-11 positive cancer cells adhere to and migrate back and forth dynamically on the cell bodies of fibroblasts. By adhering to fibroblasts for co-invasion through 3-D matrices, cancer cells acquire higher invasion speed and velocity, as well as significantly elevated invasion persistence, which are exclusive characteristics of fibroblast invasion. Silencing cadherin-11 in cancer cells or in fibroblasts, or in both, significantly decouples such physical co-invasion. Additional bioinformatics studies and PDX (patient derived xenograft) studies link such cadherin-11 mediated cancer hijacking fibroblasts to the clinical cancer progression in human such as triple-negative breast cancer patients. Further animal studies confirm cadherin-11 mediates cancer hijacking fibroblasts in vivo and promotes significant solid tumor progression and distant metastasis. Moreover, overexpression of cadherin-11 strikingly protects 4T1-luc cells from implant rejection against firefly luciferase in immunocompetent mice. Overall, our findings report and characterize the critical post-EMT event of cancer hijacking fibroblasts in cancer progression and suggest cadherin-11 can be a therapeutic target for solid tumors with stroma. Our studies hence provide significant updates on the “EMT” theory that EMT cancer cells can hijack fibroblasts to achieve full mesenchymal behaviors in vivo for efficient homing, growth, metastasis and evasion of immune surveillance. Our studies also reveal that cadherin-11 is the key molecule that helps link cancer cells to stromal fibroblasts in the “Seed & Soil” theory.

**Graphical Abstract:** 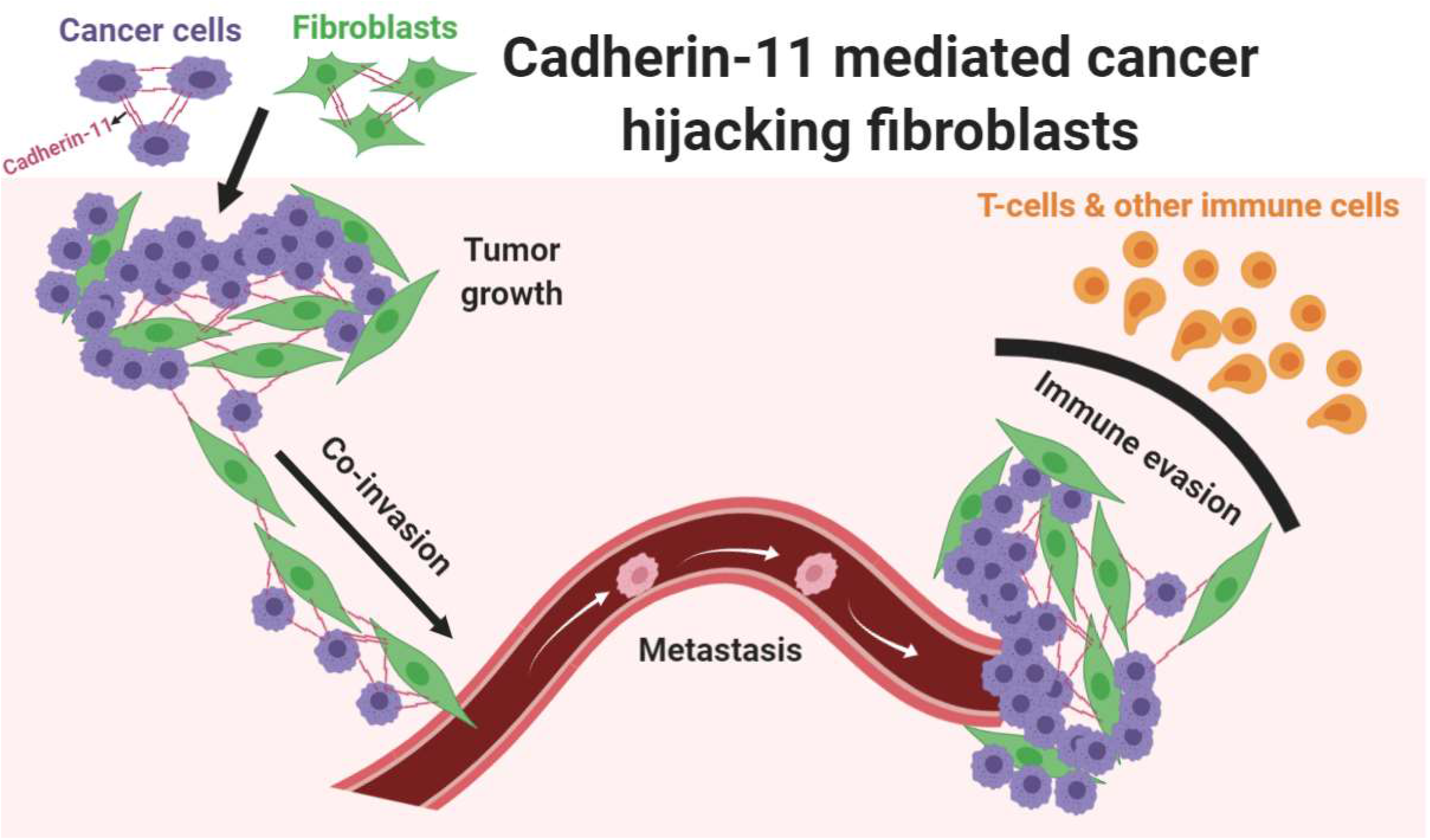

## Introduction

The prevailing concept for the metastasis of mesenchymal-like cancer cells is that they have undergone the EMT (epithelial-mesenchymal transition) process and hence have acquired the mesenchymal cell characteristics of being able to invade by themselves ^1–3^. This is evidenced by the synergistic orchestra between the EMT-TFs (EMT-activating transcription factors) signaling network and the cell migration/invasion signaling networks including integrins, MMPs (metalloproteinases), RTKs (Receptor tyrosine kinases) and small GTPases ^4,5^. EMT has also been suggested to promote primary tumor growth and progression by linking the EMT-TFs signaling to multiple growth factor signalings and cell division signaling in the cancer cells. However controversially, a partial EMT process with adaptive plasticity to the homing microenvironment and/or a complete reversal of EMT process called MET (mesenchymal-epithelial transition) might be involved or even required for successful homing and growth of some types of cancer in vivo ^6–8^. Till today, the critical role of EMT in cancer in vivo is still actively debated ^3^.

In recent years, cancer-associated fibroblasts (CAFs) have been suggested to not only support cancer growth ^9,10^ and contribute to the inflammation in the tumor microenvironment ^11–13^, but also to play an important role in local cancer invasion and cancer distal metastasis ^14–16^. For epithelial-like cancer cells, it is relatively straightforward to understand that they follow fibroblasts created micro-tunnels to invade through 3-D matrices ^17,18^ since epithelial-like cancer cells are usually not capable by themselves of invading through a dense 3-D microenvironment filled with extracellular matrices. However, for mesenchymal-like cancer cells, the story becomes much more complicated. In a solid tumor microenvironment, with both mesenchymal-like cancer cells and fibroblasts which can invade through 3-D matrices by themselves, a simple question to ask would be whether fibroblasts are facilitating or dampening the mesenchymal cancer cell invasion.

With the above question left to be answered in detail, recent studies on the co-invasion of these two mesenchymal cell types in the same 3-D microenvironment have reported that fibroblasts facilitate cancer cell invasion ^19,20^ and a direct contact between human lung cancer cells and fibroblasts is essential during their co-invasion through 3-D matrices, however the detailed mechanisms are largely unknown ^21^.

Cadherin-11, also called OB-cadherin, a type II classical cadherin, was initially discovered in osteoblasts ^22^. In the past twenty years, the Brenner lab and peers have observed that cadherin-11 is also specifically expressed in fibroblasts from various tissues ^23–26^. It has also been reported that there is a strong positive correlation between the protein expression level of cadherin-11 in breast cancer cells and the severity of their metastasis in vivo ^27^. The cadherin expression switch from E-cadherin to cadherin-11 in cancer cells has also been defined as one of the major cancer EMT markers ^28,29^. Cadherins regulate homophilic cell-to-cell adhesion, which has been classically defined as one type of cadherin binds to the same type of cadherin on the other cell in trans ^30,31^. Such cadherin adherens junction (AJ) formation happens generally in the same cell types. Interestingly, in a solid tumor microenvironment, there are cadherin-11 expressions in both mesenchymal-like cancer cells and stromal fibroblasts. Thus, it is the homophilic adhesion at the molecular level (cadherin-11 to cadherin-11), but heterophilic adhesion at the cellular level (cancer cells to fibroblasts). Hence, in this study, we focused on the critical post EMT events happening after the cadherin switch from E-cadherin to cadherin-11 in cancer cells. We scrutinized the role of cadherin-11 dependent cancer-to-fibroblast adhesion in multiple 3D ex vivo cell co-invasion models. We then determined the correlation and clinical relevance of cadherin-11 mediated cancer hijacking fibroblasts in human breast cancer progression by employing most up-to-date bioinformatics databases. We further studied how such cadherin-11 mediated cancer hijacking fibroblasts post EMT events contribute to cancer growth, local invasion, distant metastasis, progression and immune rejection against tumor implants in multiple mouse models in vivo.

## Results

### Cadherin-11 positive cancer cells form adherens junctions (AJs) with fibroblasts

The adherens junction has been defined as a cell-cell junction where the cadherins on one cell bind to the same type of cadherins on the other cell (homophilic binding). Hence, it has been largely studied in cultures of a single cell type ^30,31^. Adherens junction formation also requires a whole set of intracellular AJ structural proteins, such as catenins, to stabilize such cell-to-cell cadherin binding. It is not known if the cancer cells that express cadherin-11 have enough molecular machinery to be capable of forming stable AJs with fibroblasts that endogenously express abundant cadherin-11 proteins ^26,32^. To address this question, we first cocultured MDA-MB-231 or BT549 breast cancer cells, which inherently express cadherin-11 proteins ^27^, with primary human fibroblasts and stained for cadherin-11 protein. The results confirmed cadherin-11 AJ formation between these human cancer cells and human fibroblasts (Figure 1A & B).

**Figure 1.**
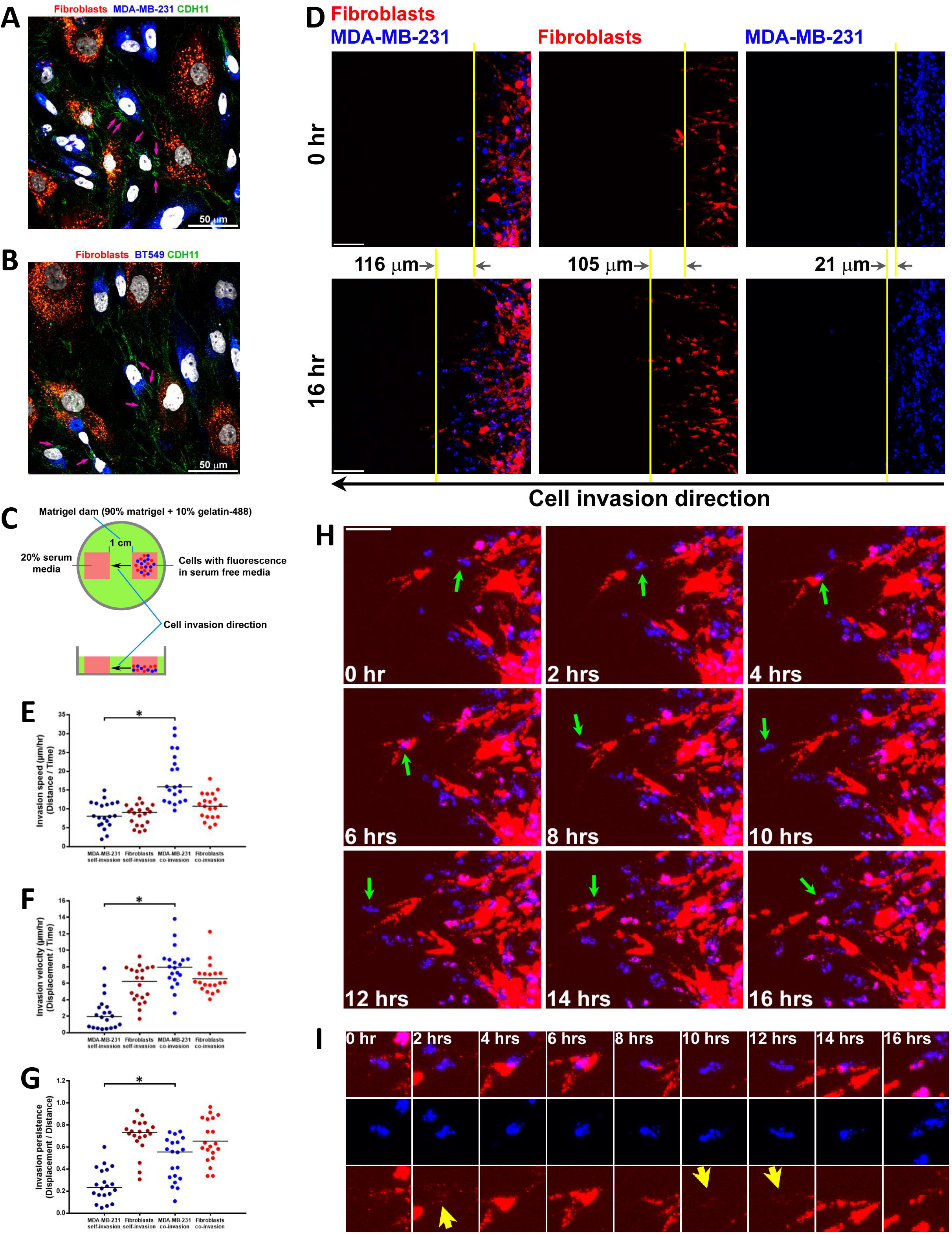
Cancer cells hijack fibroblast cell bodies for invasion through 3D matrices in the matrigel dam assay. A & B. Primary human fibroblasts were pre-labeled by DiI (red) and MDA-MB-231 (A) or BT549 cells (B) were pre-labeled by DiD (blue). Cells were cocultured for overnight before fixation for IF staining. Cadherin-11 protein was stained by the specific monoclonal primary antibody (clone 16A) followed by the secondary antibody staining (green). Purple arrows are pointing to the cadherin-11 AJs between cancer cells and fibroblasts. Confocal Z-stacks were scanned from the top of the cell to the bottom of the cell. All 2-D images shown were from 3-D Z-stacks maximum projections. C. The schematic to describe the cell invasion assay. D. Fibroblasts and MDA-MB-231 cells were pre-labeled as before. Fibroblasts alone (middle panels), MDA-MB-231 alone (right panels) or Fibroblasts and MDA-MB-231 (left panels) together in a 1:1 ratio were subjected to the cell invasion assay as described in (C). The whole cell population invasion displacements on the X-axis with a direction to the left were labeled between the yellow lines of 0 hr and 16 hr. The green fluorescence from the matrigel was omitted to clearly visualize the cells. Size bar, 100 μm. E, F & G. Cell invasion speed (Distance/time), invasion velocity (Displacement on the X-axis/time) and invasion persistence (Displacement/Distance) were quantitated. *, P < 0.05. H. Time-lapse zoom-in panels from Video 3. Green arrows are pointing at one cancer cell that was invading back-n-forth by attaching to and sliding on the cell bodies of fibroblasts. The red channel imaging offsets were elevated to visualize the long but thin invasive protrusions of fibroblasts. Size bar, 100 μm. I. Cropped and zoom-in panels from (H) to present the details of the invasive protrusions of fibroblasts (yellow arrow heads).

### Cancer cells hijack fibroblast cell bodies for invasion through 3D matrices

Next, to observe the cell co-invasion capabilities of these cancer cells and fibroblasts, we developed a “matrigel dam” directional 3D cell invasion assay in which we can adjust the serum concentration or the matrigel dam width to control cell invasion speed (Figure 1C). We determined that 20% serum driving cell invasion through a 1 cm wide matrigel dam showed a significant invasion speed difference between fibroblasts and cancers during their individual invasion in this assay. Pure fibroblasts (Figure 1D, middle panels) invaded much faster with much higher persistence than the pure cancer cells (Figure 1D, right panels) under the same conditions (Figure 1E, F & G). However, strikingly, when fibroblasts and cancer cells were mixed together (1:1 ratio), these cancer cells physically associated with the fibroblasts and they managed to invade faster and also as persistently as the fibroblasts in this assay (Figure 1D, left panels, Figure 1E, F & G, Video 1, 2 & 3), like taking a high-speed train for faster travel.

High magnification zoom-in confocal imaging with elevated offsets on the red fluorescence channel revealed that cancer cells physically stayed adhered to the fibroblasts and migrated back and forth on the fibroblast body with high dynamics when these fibroblasts invaded through the 3-D matrix (Figure 1H & I, Video 4). We also observed that fibroblasts formed long and thin cell body protrusions at the invasion front to invade through the 3-D matrix and cancer cells remained physically adhered to these long and thin fibroblast protrusions throughout the co-invasion (Figure 1I). In such cancer cell-fibroblast co-invasion, almost none of the cancer cells invaded without adhering to fibroblasts (Figure 1H & I, Video 4). Multiphoton microscopy imaging of a large cell population of cancer cells and fibroblasts during their co-invasion through 3D matrices in another cell invasion assay also confirmed strong physical adhesions between the cancer cells and the fibroblasts during their co-invasion (Figure S1A &B).

To avoid the possible concerns of cells touching the glass surface beneath the matrigel dam during invasion, a possible mixed 2.5-D ^33^ and 3-D cell invasion (Figure 1C), we employed a full 3-D spheroid cell invasion assay (Figure 2A) ^34^. Interestingly in such assays, pure cancer cells expressing cadherin-11 did not invade much out of the spheroids into the surrounding 3-D matrices. By contrast, we observed significant cell invasion in all the spheroids that were composed of pure fibroblasts (Figure 2B, upper panels, and Figure 2C). When these cancer cells were mixed with fibroblasts in a 1:1 ratio (total cell numbers were maintained the same in all spheroids), we again observed striking cancer cell invasion together with the fibroblasts. Further, the cell bodies of many cancer cells partially co-localize with the cell bodies of the leading fibroblasts (Figure 2B, bottom panels and zoom panels), confirming a proximal physical contact between cancer cells and fibroblasts during their co-invasion, as previously observed in our live cell imaging (Figure 1H & I, Video 4).

**Figure 2.**
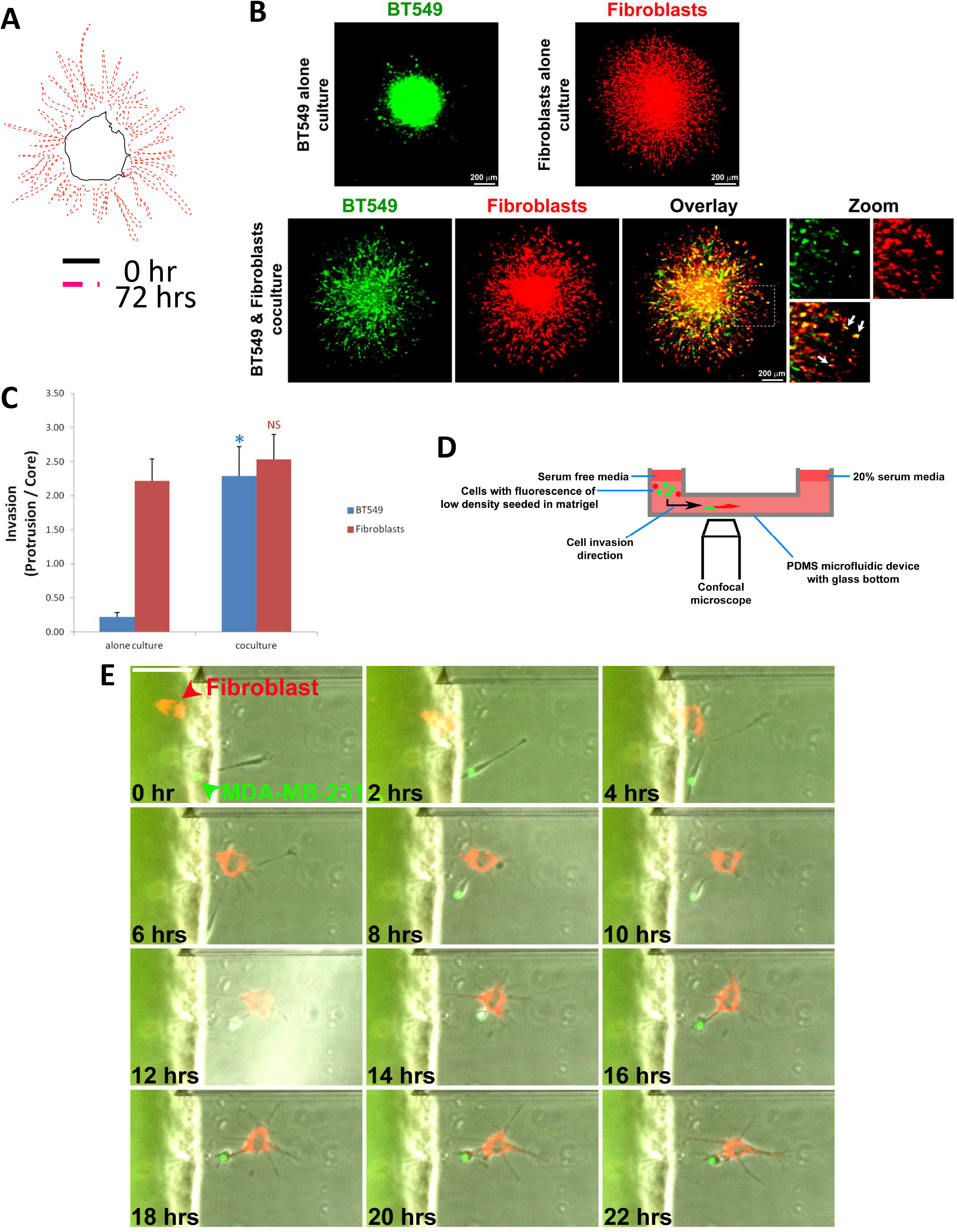
Cancer cells hijack fibroblast cell bodies for invasion through 3D matrices in the spheroid assay and the microfluidic assay. A. The schematic explaining the invasion quantitation for the spheroid invasion assay. Invasion ratio was calculated by the area of the protrusion divided by the area of the core. B. Primary human fibroblasts were pre-labeled by DiI (red) and BT549 cells were pre-labeled by DiO (green). BT549 alone spheroid (top left), fibroblasts alone spheroid (top right), or BT549 and fibroblasts coculture (1:1) spheroid (bottom panels) were subjected to the 3D spheroid cell invasion assay. Total cell numbers maintained the same in every spheroid. Confocal Z-stacks were scanned from the top of the spheroid to the bottom. All 2-D images shown were from 3-D Z-stacks maximum projections. C. Quantitation for images as in (B). Experiments were repeated 6 times (n=6). *, P < 0.05. D. The schematic describing the microfluidic invasion assay. E. Time-lapse panels from Video 5. Primary human fibroblasts were pre-labeled by DiI (red) and MDA-MB-231 cells were pre-labeled by DiO (green).

Next, we sought to confirm such cancer cell and fibroblast co-invasion at the single-cell level without forcing proximal physical contacts by mixing thousands of cells in high densities to begin with as in the previous two cell invasion assays. By employing a microfluidic cell invasion device that is able to limit flow to allow just a couple of cells to pass through a narrow tunnel filled with 3-D matrices (Figure 2D) ^35^, we observed such “cell hijacking” between one single cancer cell and one single fibroblast. It is important to point out that both cells started their own invasion as two separated single cells without any contact and such “cell hijacking” happened at the first contact as soon as they touched each other (Figure 2E & Video 5). Interestingly, when the cancer cell was caught by the fibroblast, the cancer cell body almost totally rounded up while the fibroblast still maintained its invasive cell shape in the 3-D matrices. This suggests the cancer cell gave up its self-invasion but took the “train” (fibroblast) to travel. Thus, it is reasonable to describe such co-invasion as the cancer cell “hijacking” the fibroblast, but not vice versa (Video 5).

### Co-invasion by cancer cells hijacking fibroblasts is cadherin-11 dependent

As cadherin-11 AJs can form between these cancer cells and fibroblasts (Figure 1A & B), we next sought to determine the role of cadherin-11 in regulating such cancer cells hijacking fibroblasts for co-invasion. First, we confirmed that cadherin-11 siRNA successfully downregulated cadherin-11 protein expression in these cells (Figure S2A & B). In the same 3-D spheroid cell invasion assay as described previously, we observed that cadherin-11 downregulation promoted cell dissemination in these breast cancer cells (Figure 3A) when they were cultured alone, suggesting that same cell adhesion suppresses invasion of these cancer cells. In contrast, cadherin-11 downregulation inhibited cell invasion in a culture of sole fibroblasts (Figure 3B). Interestingly, we observed totally opposite cell invasion behaviors by equivalently perturbing cadherin-11 in human cancer cells and human primary fibroblasts (Figure 3A & B), which is not a too big surprise for us according to our previous publications about the clear differences between cancer cells and normal fibroblasts ^36^. Cell-to-cell adhesion forces in AJs and TJs (tight junctions) are usually defined as “frictions” during cell dissemination or cell invasion ^37,38^. Hence, it is easy to understand that when cell-to-cell adhesion forces in AJs are weakened by cadherin-11 downregulation, cell dissemination increases in these breast cancer cells. However, our results in fibroblasts from the same spheroid assay suggest that the “cell-to-cell adhesion friction” theory is not enough to explain the phenomenon in fibroblasts. Interestingly, this finding is consistent with previous studies, involving several different 3-D assays which reported that cell invasion significantly decreases in fibroblasts with a cadherin-11 blockade by a cadherin-11-Fc fusion protein or in fibroblasts from cadherin-11-null mice ^32,39^. Also, according to our findings above in untransfected wildtype cells, cancer cells alone didn’t invade much while fibroblasts alone did invade a lot in the 3-D spheroid assay, suggesting that fibroblasts do have a strong driving power for invasion but cancer cells don’t in this case (Figure 2B). Thus, by integrating information from all these studies, we see that, in contrast to the effects of cadherin-11 friction forces in regulating cell invasion, there exists an unknown mechanism by which cadherin-11 actually promotes cell invasion in fibroblasts. But such a mechanism could be negligible or nonexistent in all the cancer cells that we have tested (Figure S2C).

**Figure 3.**
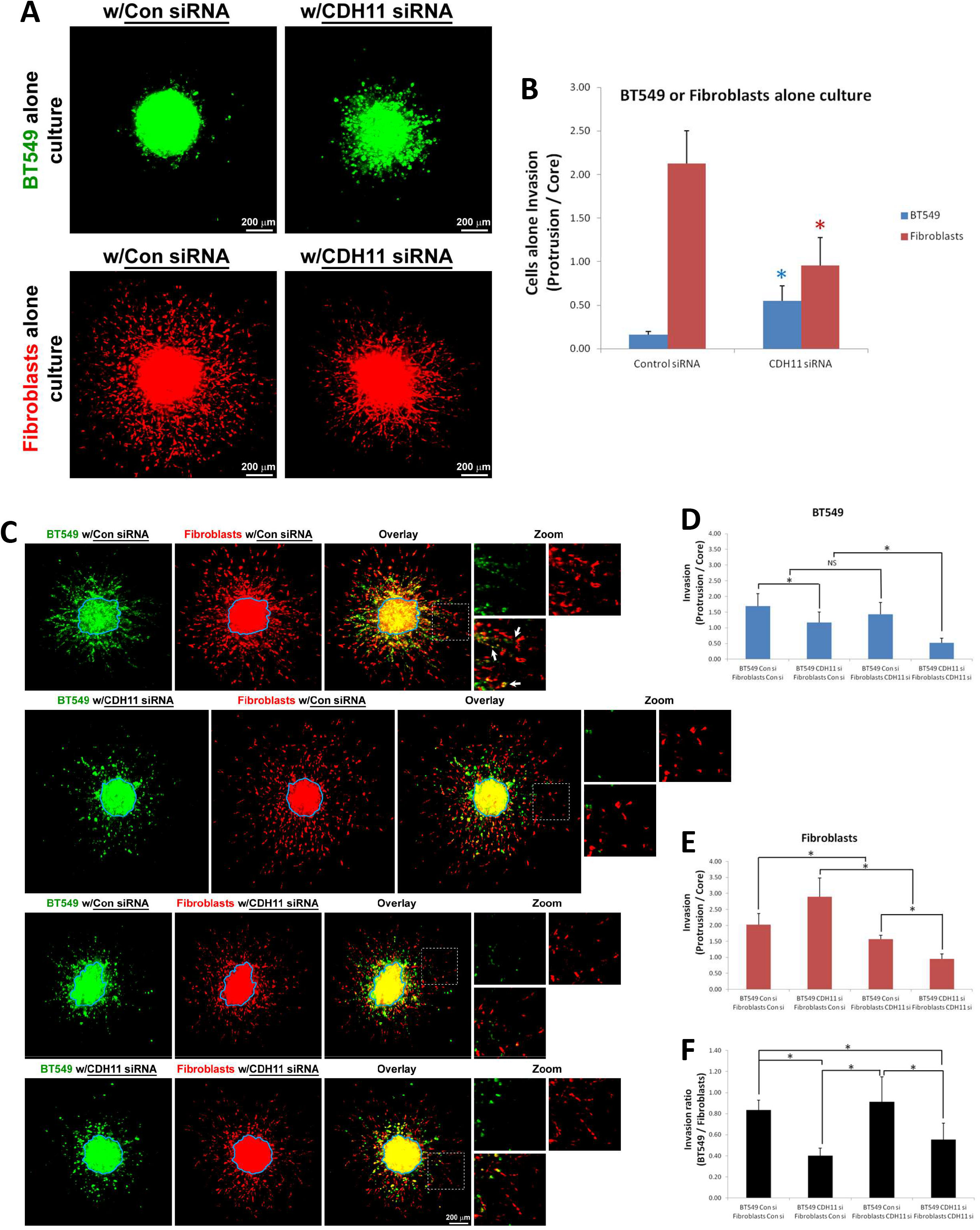
The cancer cell hijacking fibroblast invasion is cadherin-11 dependent. A. Primary human fibroblasts were pre-labeled by DiI (red) and BT549 cells were pre-labeled by DiO (green). BT549 alone spheroid with negative control siRNA or CDH11 siRNA (top panels), and fibroblasts alone spheroid with negative control siRNA or CDH11 siRNA (bottom panels) were subjected to the 3D spheroid cell invasion assay. B. Quantitation for images as in (A). Experiments were repeated 6 times (n=6). *, P < 0.05. C. BT549 and fibroblasts coculture (1:1) spheroid with negative control siRNA or CDH11 siRNA were subjected to the 3D spheroid cell invasion assay. D, E & F. Quantitation for images as in (C). Experiments were repeated 6 times (n=6). *, P < 0.05. Total cell numbers maintained the same in every spheroid. Confocal Z-stacks were scanned from the top of the spheroid to the bottom. All 2-D images shown were from 3-D Z-stacks maximum projections.

Hence, based on these findings, we speculate that fibroblasts are more like the “driving wheel” in a car where cadherin cell-to-cell adhesion friction plays as the resistance force and the pulling force between each adjacent cell concurrently. Whilst in cancer cells, which are more like the “non-driving wheel” in a car, cadherin cell-to-cell adhesion friction most likely only plays as the resistance force because the cancer cells themselves don’t have a potent power for invasion by themselves in the 3-D assay (Figure 2B BT549 alone culture panel; Figure 3A BT549 alone culture w/Con siRNA panel). However, in fibroblast and cancer cell coculture, fibroblasts are able to transmit power to drive the cancer cells to invade (Figure 2B coculture panel). Next, we started to perturb cadherin-11 in the coculture system by silencing cadherin-11 in cancer cells only, or in fibroblasts only, or in both concurrently. Results revealed that in all cadherin-11 silencing cases, the co-invasion of cancer cells hijacking fibroblasts was significantly de-coupled (Figure 3C-F; Figure S3A-D). Interestingly, silencing cadherin-11 in just cancer cells significantly drove fibroblasts to invade farther in the 3D coculture system (Figure 3C, second row), which further suggests that cadherin cell-to-cell adhesion friction in cancer cells plays as the resistance force. When such resistance force is reduced by cadherin-11 silencing in these cancer cells, the neighboring fibroblasts were freed and let go for invasion. Thus, all these results suggest that, in coculture, fibroblasts transmit power to drive cancer cell invasion and such power transmission (most likely as the force transmission by physical pulling) is cadherin-11 cell-to-cell adhesion dependent.

### Cadherin-11 is specifically expressed in triple negative breast cancer cells (TNBCs) and its high expression is associated with poor clinical outcome in breast cancer patients

Bioinformatics analysis based on the CCLE (Cancer Cell Line Encyclopedia) ^40,41^ database reveals that cadherin-11 is highly expressed in a subset of triple negative breast cancer cells (TNBCs), but not non-TNBCs (Figure 4A).

**Figure 4.**
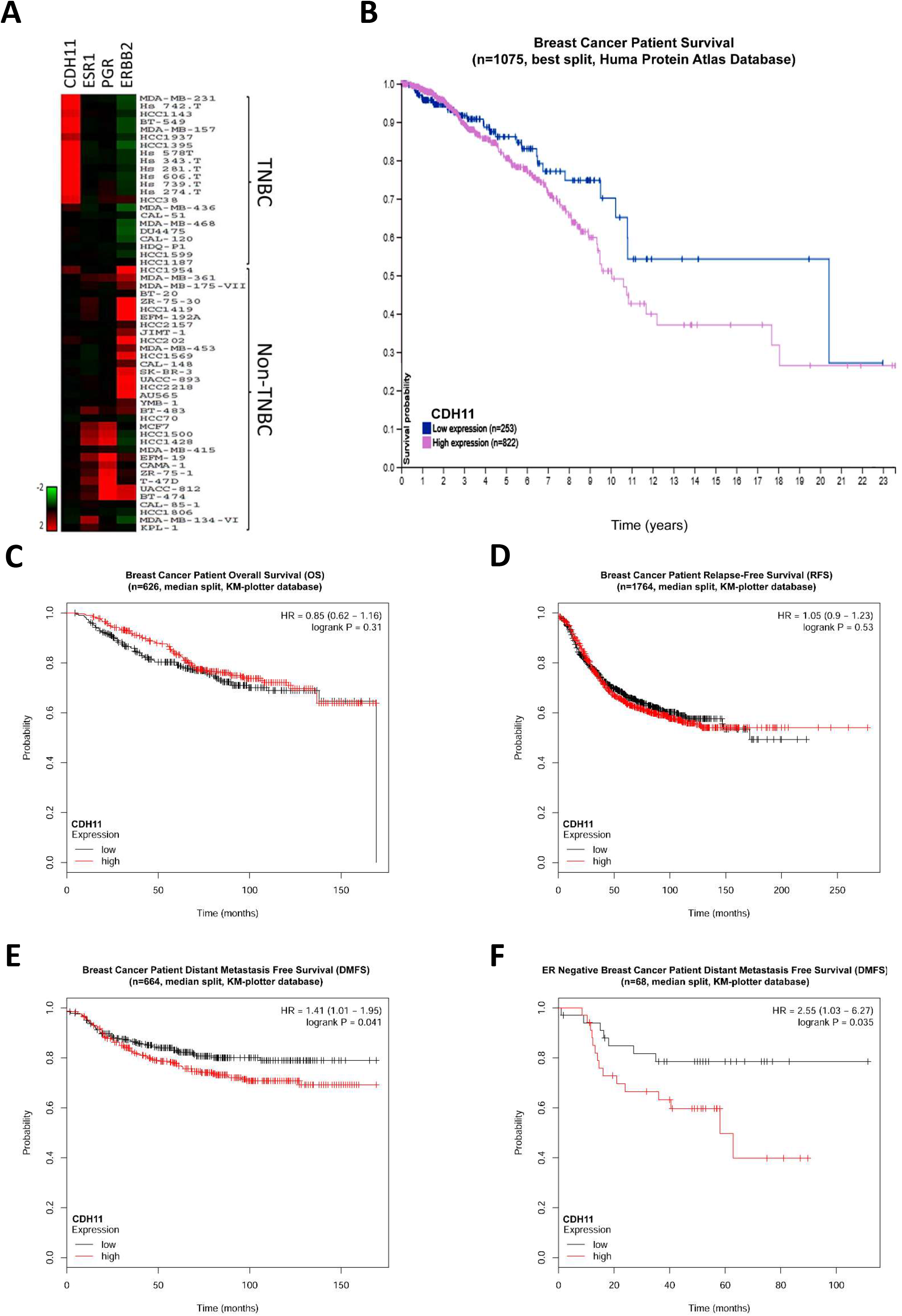
Cadherin-11 is specifically expressed in triple negative breast cancer cells (TNBCs) and its high expression is associated with poor clinical outcome in breast cancer patients. A. CDH11 (Cadherin-11), ESR1(Estrogen Receptor 1), PGR (Progesterone Receptor) & ERBB2 (Receptor tyrosine-protein kinase erbB-2) expression data in breast cancer cell lines from CCLE (Cancer Cell Line Encyclopedia, The Broad Institute of MIT & Harvard) were plotted into a heat map. B. Kaplan-Meier analysis for breast cancer patients stratified by CDH11 expression for 1075 patients from The Human Protein Atlas database. C-E. Kaplan-Meier plots of overall survival (C, n=626), relapse-free survival (D, n=1764) and distant metastasis free survival (E, n=664) of breast cancer patients in relation to CDH11 expression according to The KM-plotter database. F. Kaplan-Meier plots of distant metastasis free survival (n=68) of ER negative breast cancer patients in relation to CDH11 expression according to The KM-plotter database.

Clinical data from The Pathology Atlas of The Human Protein Atlas database (www.proteinatlas.org) ^42^ shows high expression of cadherin-11 correlates with poor patient survival in 1,075 breast cancer patients (Figure 4B). Next, we analyzed the clinical data from the breast cancer mRNA gene chip dataset of the KM-plotter database (www.kmplot.com/analysis/) ^43^. Interestingly, high expression of cadherin-11 doesn’t correlate with poor patient survival either in 626 patients by using the Overall Survival (OS) analysis method (Figure 4C) or in 1,764 patients by using the Patient Relapse-Free Survival (RFS) analysis method (Figure 4D). Instead, high expression of cadherin-11 significantly (logrank P = 0.041) correlates with poor patient survival in 664 patients by using the Distant Metastasis Free Survival (DMFS) analysis method (Figure 4E). Such an interesting finding from the patient survival clinical data clearly positions the significant role of cadherin-11 in cancer metastasis, which links up with our findings of cancer hijacking fibroblasts for invasion in 3D ex vivo studies above (Figures 1, 2 & 3). Next, we sought to analyze the correlation between high expression of cadherin-11 with patient survival in triple negative breast cancer patients by using the DMFS analysis method as in Figure 4E. However, due to the very limited number of triple negative breast cancer patient survival data, instead, we analyzed the correlation between high expression of cadherin-11 with ER negative breast cancer patient survival in 68 patients by using the DMFS analysis method. Results show high expression of cadherin-11 extremely (logrank P = 0.035) correlates with poor patient survival in these patients (Figure 4F). Such clinical data further validates our findings of high cadherin-11 expression particularly in triple negative breast cancers from the CCLE database analysis as above (Figure 4A). Overall, all the clinical data from these two databases suggests high expression of cadherin-11 in cancer cells, so that they can readily hijack fibroblasts by such a cadherin-11-to-cadherin-11 homophilic adhesion at the molecular level, significantly drives cancer progression in human patients, particularly with high relevance to distant metastasis progression.

### Cadherin-11 mediates fibroblasts regulated tumor homing, growth and local invasion in human triple negative breast cancer with human fibroblast co-implantation xenograft mouse models

Optimal cadherin based adherens junction formation is achieved by one type of cadherin binds to the “same” type of cadherin on the other cell in trans ^30,31^. Previous studies on cadherin-11 mediated cancer growth and progression in xenograft mouse models were done by implanting human breast cancer cells (with “human” cadherin-11) into the mouse stroma ^44,45^. It needs to be pointed out that mouse stromal fibroblasts express “mouse” cadherin-11 but not “human” cadherin-11. Although such a mouse model can be used to test the cadherin-11 intracellular signaling changes in these cadherin-11 bearing cancer cells themselves supported by a “mouse” stromal microenvironment in vivo, it is not suitable for studying the effects of homophilic molecular adhesions of “human” cadherin-11 on the cancer cells to “human” cadherin-11 on the stromal fibroblasts, which actually happens in human.

To focus on homophilic “human” cadherin-11 to “human” cadherin-11 adhesions between cancer cells and stromal fibroblasts, we designed a human triple negative breast cancer with primary human fibroblast co-implantation xenograft mouse model, which is similar to previous in vivo mouse studies done by human cancer and human fibroblast co-implantations ^46–48^. We found that co-implantation with primary human fibroblasts significantly promoted human cancer cell growth in NOD/SCID mice as visualized by weekly bioluminescence imaging of MDA-MB-231-luc cells, especially at early time points after implantation (Figure 5A & B, note only MDA-MB-231-luc cells have bioluminescence but fibroblasts don’t). Such a finding was not obvious at later time points possibly because later cancer growth was largely supported by the mouse stroma. Further, co-implantation with primary human fibroblasts significantly raised the successful rate of MDA-MB-231-luc xenograft growth in these mice (Figure 5A, B & C, note those small tumors with bioluminescence below 1 × 10^8^ photons/s at week 11). As a control, implantation of primary human fibroblasts alone into NOD/SCID mice did not form any tumors (data not shown). Interestingly, in vivo bioluminescence imaging of MDA-MB-231-luc cells revealed that cancer cells with fibroblast co-implantation underwent more local invasion (measured as total tumor area) compared to cancer cells without fibroblast co-implantation (Figure 5D), consistent with the cell invasion findings in the ex vivo 3D spheroid assays. Tumor volume measurements further showed that tumors of MDA-MB-231-luc with fibroblast co-implantation grew much faster and larger than tumors of MDA-MB-231-luc sole implantation (Figure 5E). Such a finding was also observed in triple negative breast cancer patient derived cancer (ID: PDX5993) with or without primary human fibroblast co-implantation xenografts in NOG mice. (Figure 5F). Overall, these results confirm that co-implantation with primary human fibroblasts promote human cadherin-11 positive triple negative breast cancer cell homing, growth and local invasion in mouse in vivo.

**Figure 5.**
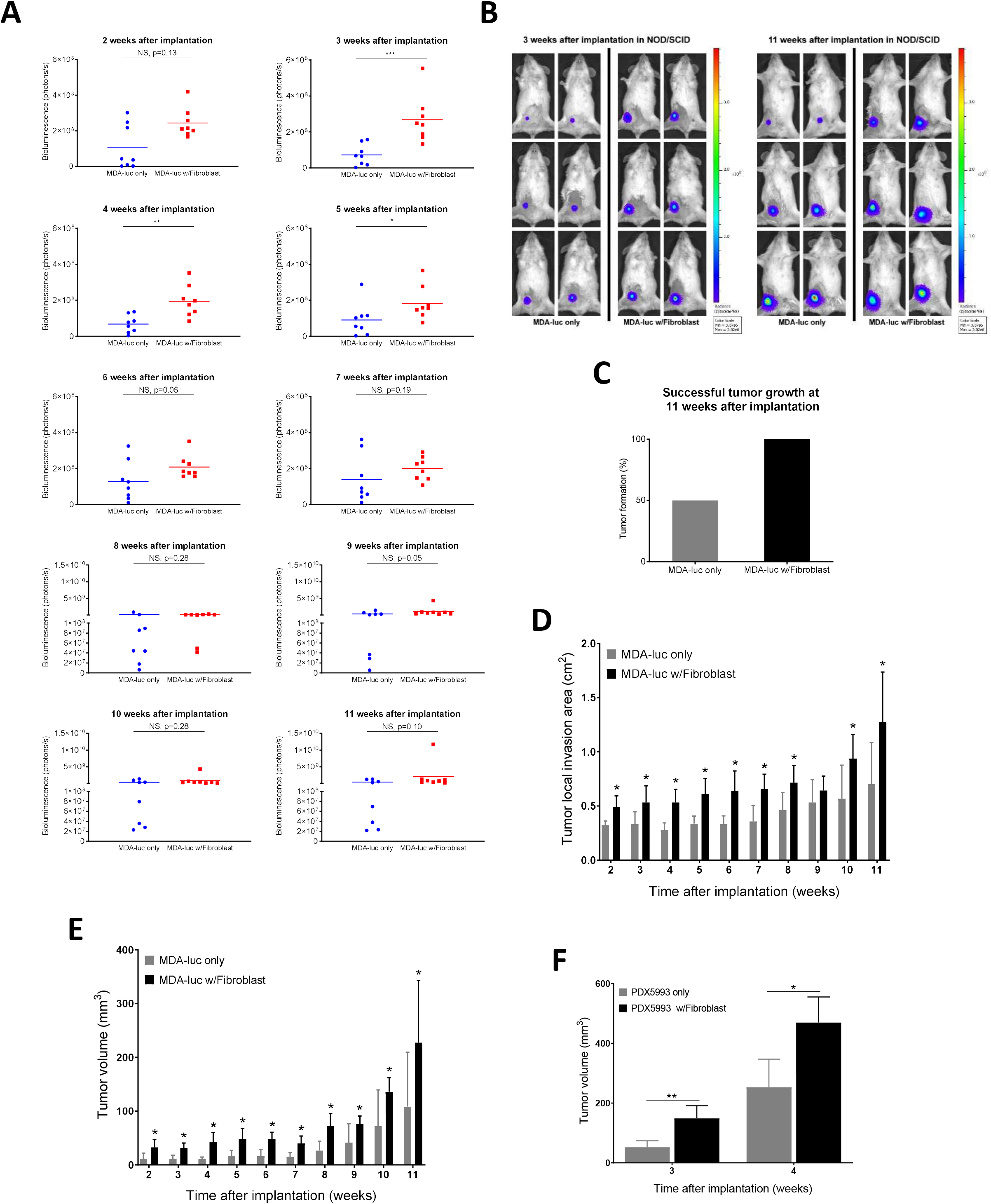
Fibroblasts regulate tumor homing, growth and local invasion in triple negative breast cancer with fibroblast co-implantation xenograft mouse models. 1 × 10^6^ of cancer cells with/without 1 × 10^6^ of primary human fibroblast cells were implanted into the left fourth mammary fat pad of NOD/SCID (for MDA-MB-231, 5A-5E), or NOG (for PDX5993, 5F) mice. A. Comparison of bioluminescence intensity in NOD/SCID mice at 2 to 11 weeks after implantation between the MDA-MB-231-Luc cells only implantation group (MDA-luc only, blue) and the MDA-MB-231-Luc cells with primary human fibroblasts co-implantation group (MDA-luc w/Fibroblast, red). B. Representative *in vivo* bioluminescence images from (A) in the early stage (3 weeks, left panel) and the late stage (11 weeks, right panel) post implantation. C. Fibroblasts supported tumor formation at 11 weeks after implantation. Tumors with total bioluminescence above 1 × 10^8^ photons/s were considered as successful growth. D. Fibroblasts stimulated tumor local invasion as quantified by the total area of the visualized bioluminescence from MDA-MB-231-luc cells in vivo with or without fibroblast co-implantation. E. Comparison of tumor volume in NOD/SCID mice at 2 to 11 weeks after implantation between the MDA-MB-231-Luc cells only implantation group and the MDA-MB-231-Luc cells with primary human fibroblasts co-implantation group. F. Comparison of tumor volume in NOG mice at 3 and 4 weeks after implantation of patient-derived triple negative breast cancer cell xenograft PDX5993 with or without primary human fibroblast co-implantation. Data were presented as mean ± SD. n=8 in (A-E). n=4 in (F). P values were determined by two-tailed Student’s t tests (NS, not significant; *, 0.01 < p < 0.05; **, p < 0.01).

Previous studies on cadherin-11 in cancer progression in vivo were performed mainly by perturbing cadherin-11 expression level in cancer cells only to determine how cadherin-11 regulates cancer cells themselves, including studies on cadherin-11 regulated intracellular signalings and cadherin-11-to-cadherin-11 homophilic adhesions between the same type of cancer cells ^44,45^. To avoid the effects of perturbing cadherin-11 in altering cancer cells themselves and to focus on how the cadherin-11-to-cadherin-11 adhesions between the two different types of cells regulate human cancer hijacking human stromal fibroblasts in vivo, we chose to perturb cadherin-11 only in primary human fibroblasts in our human triple negative breast cancer with primary human fibroblast co-implantation xenograft mouse model. We found that the reduction of cadherin-11 expression in primary human fibroblasts (Figure 6A) significantly reduced the enhancement of MDA-MB-231-luc growth in vivo (Figure 6B & C) as well as the whole tumor growth (Figure 6D). Overall, these results confirm that human cancer hijacking human stromal fibroblasts in vivo with enhanced cancer progression is cadherin-11-to-cadherin-11 adhesion dependent.

**Figure 6.**
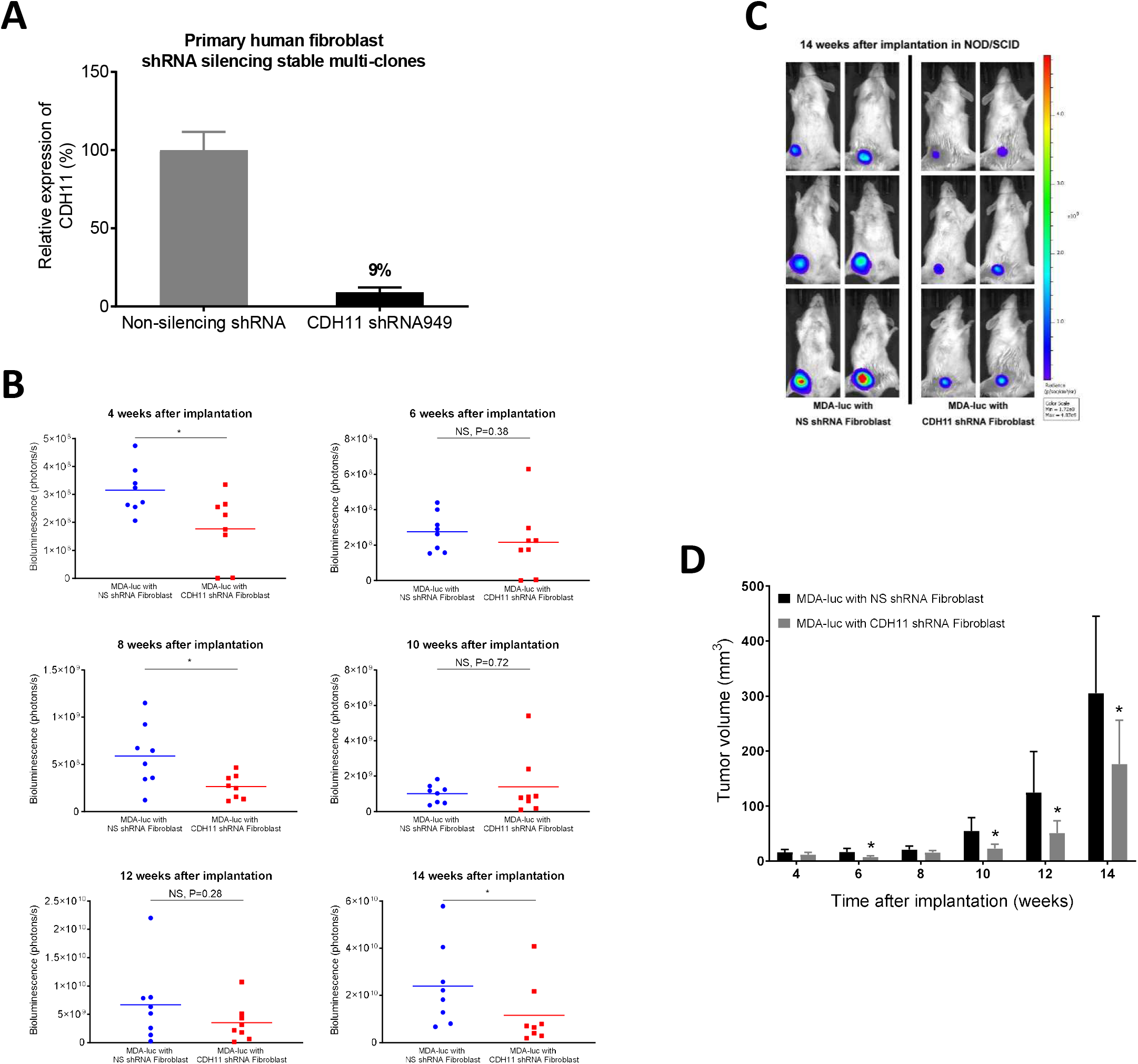
Silencing of cadherin-11 in primary human fibroblasts suppresses tumor progression in the triple negative breast cancer with fibroblast co-implantation xenograft mouse model. A. Primary human fibroblasts were transduced by GIPZ lentiviral CDH11 shRNA or non-silencing shRNA with a GFP reporter. Stable transductant cells were sorted out by FACS based on the GFP signal. Silencing of CDH11 in these cells was then quantified by RT-qPCR. B. Effect of CDH11 silencing in human fibroblasts on cancer cell growth in the cancer with fibroblast co-implantation xenograft mouse model. 1 × 10^6^ of MDA-MB-231-luc cells mixed with 1 × 10^6^ of stable non-silencing (blue) or CHD11 silencing (red) primary human fibroblasts were co-implanted into the left fourth mammary fat pad of NOD/SCID mice. The bioluminescence of the MDA-MB-231-Luc cells were measured every 2 weeks. C. Representative *in vivo* bioluminescence images from (B) at 14 weeks after cancer with fibroblast co-implantation. D. Comparison of tumor volume in mice co-implanted with MDA-MB-231-luc cells and non-silencing or CDH11 silencing primary human fibroblasts. Data were presented as mean ± SD (n=8). P values were determined by two-tailed Student’s t tests (NS, not significant; *, 0.01 < p < 0.05).

### Overexpression of cadherin-11 in 4T1 mouse triple negative breast cancer cells to enhance their adhesion to mouse fibroblasts promotes tumor invasion, growth and distal metastasis

Next, we sought to determine the in vivo effects of cadherin-11 overexpression in cancer cells with their enhanced adhesion to stromal fibroblasts. 4T1 murine cancer implantation in BALB/c mice is a widely used immunocompetent triple negative breast cancer implantation mouse model, which very closely mimics triple negative breast cancer progression in human ^49,50^. Interestingly, although 4T1 cells are very invasive breast cancer cells, they express high levels of epithelial E-cadherin with almost no expression of mesenchymal N-cadherin or cadherin-11 (Figure S4A) ^51^. Thus, we firstly made stable cadherin-11 overexpressing 4T1 cell lines and confirmed the overexpression levels by Real-Time qRT-PCR (Figure 7A). Next, in our established ex vivo 3D spheroid assay, we observed that unlike the human BT549 TNBC cell line that showed little invasion on its own (Figure 2B), 4T1 wildtype cells showed invasion which was not enhanced by cadherin-11 overexpression (Figure 7B, upper panels, and Figure 7C). Furthermore, addition of mouse NIH3T3 fibroblasts did not enhance the invasion of 4T1 wildtype cells (Figure 7B & C). NIH3T3 fibroblasts did enhance the invasion of cadherin-11 overexpressing 4T1 cells (Figure 7B & C). In vivo experiments of 4T1 implantation into the mammary pads of BALB/c mice determined that tumors of 4T1 cells with mouse cadherin-11 overexpression grew significantly faster than those of 4T1 wildtype cells (Figure 7D). This is not due to an increased proliferation rate (Figure 7E). Moreover, BALB/c mice carrying tumors of 4T1 cells with mouse cadherin-11 overexpression died much earlier than those carrying tumors of 4T1 wildtype cells (Figure 7F). Micro-MRI imaging detected large metastatic tumor lesions at multiple lymph nodes under the dorsal neck skin in BALB/c mice carrying tumors of 4T1 cells with mouse cadherin-11 overexpression at the left 4^th^ mammary pads. In comparison, Micro-MRI imaging failed to detect any metastatic tumor lesion in BALB/c mice carrying tumors of 4T1 wildtype cells (Figure 7G). Overall, all these results suggest that 4T1 TNBC cells with mouse cadherin-11 overexpression can hijack mouse fibroblasts to promote tumor homing, growth, progression and metastasis with much shorter mouse survival in vivo.

**Figure 7.**
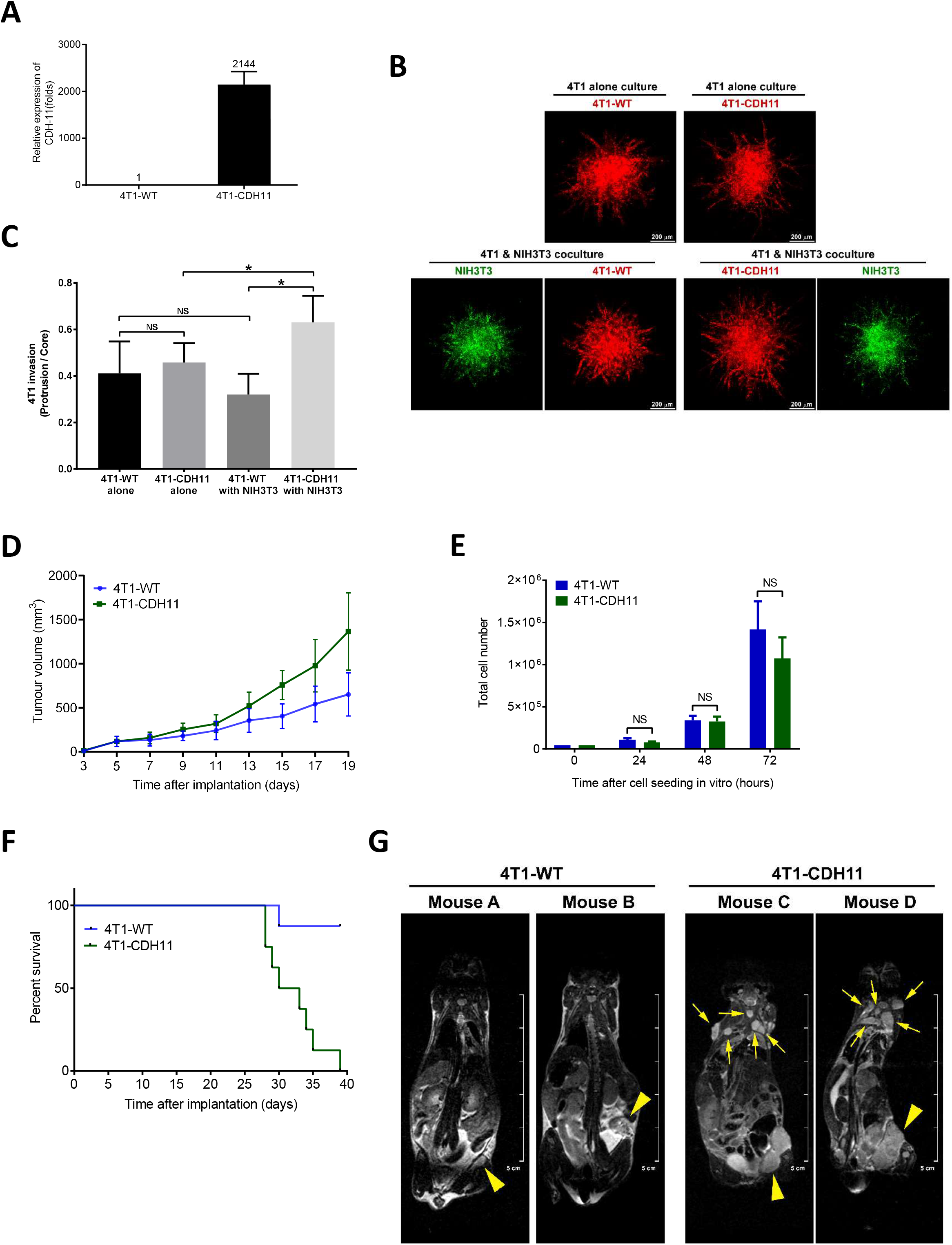
Overexpression of cadherin-11 in 4T1 mouse triple negative breast cancer cells to enhance their adhesions to mouse fibroblasts promotes tumor invasion, growth and distal metastasis. A. CDH11 stable overexpression in 4T1 cells (4T1-CDH11) was quantified by RT-qPCR against CDH11 expression levels in 4T1 wildtype cells (4T1-WT). B. 4T1-WT cells or 4T1-CDH11 cells were pre-labeled by DiD (red). NIH3T3 mouse fibroblasts were pre-labeled by DiO (green). 4T1 alone spheroids (top panels), or 4T1 and NIH3T3 coculture (1:1) spheroids (bottom panels) were subjected to the 3D spheroid cell invasion assay as in Figure 2B. Total cell numbers maintained the same in every spheroid. Confocal Z-stacks were scanned from the top of the spheroid to the bottom. All 2-D images shown were from 3-D Z-stacks maximum projections. C. Quantitation for images as in (B). Experiments were repeated 5 times (n=5). NS, not significant; *, P < 0.05. D. Comparison of tumor volume in BALB/c mice implanted with 4T1-WT cells or 4T1-CDH11 cells. 1 × 10^6^ of 4T1 cells were implanted into the left fourth mammary fat pad in BALB/c mice. Data were presented as mean ± SD (n=8). E. Comparison of 4T1-WT cell and 4T1-CDH11 cell proliferation in 2D culture in vitro. Same number of cells (46,000 cells) of each group were seeded in one well of a 6-well plate. Cell number was counted at 24 hrs, 48 hrs & 72 hrs (n=3 for each cell group at each time point) after cell seeding. NS, not significant. Note all cells were still not confluent at 72 hrs in each well of a 6-well plate. F. Kaplan-Meier survival curve of BALB/c mice implanted with either 4T1-WT cells or 4T1-CDH11 cells as in (D), n=8. G. micro-MRI imaging of tumor-bearing BALB/c mice from (D) on the 21^st^ day after implantation of 4T1-WT cells or 4T1-CDH11 cells. Multiple distal metastatic sites in the dorsal neck region lymph nodes (denoted by small arrows) were detected in mice with 4T1-CDH11 tumors. Large arrowheads denote the original tumors at the left fourth mammary fat pad. micro-MRI image Z-stacks were scanned from the dorsal side to the ventral side of the mice. Single plane image section across the dorsal neck region lymph nodes from two representative mice from each group is shown. No distal metastasis was detected in any mice with 4T1-WT tumors in all micro-MRI image Z-stacks.

### Overexpression of cadherin-11 in 4T1 mouse triple negative breast cancer cells with firefly luciferase prevents immune rejection in immunocompetent BALB/c mice in vivo

Recent studies discovered that implantation of 4T1-luc cells into immunocompetent BALB/c mice induces immune rejection against the tumor implant by the mouse immune system through recognition of the firefly luciferase protein as a foreign antigen ^52^. Such a phenomenon is also described on the product sheet of the commercial 4T1-luc cells (PerkinElmer Inc). Initially, we planned the experiments of implanting 4T1-luc cells into BALB/c mice to determine if we could achieve a mild immune rejection so that we were able to observe 4T1-luc cell activities in vivo by bioluminescence imaging. However, a very strong immune rejection against the implanted wildtype 4T1-luc cells was observed. Very unexpectedly, in comparison, a much milder immune rejection was observed in the 4T1-luc cells with stable mouse cadherin-11 overexpression (Figure 8A, B & C). To further confirm such an interesting and unexpected finding, we repeated the in vivo mouse experiments with different batches of cells for implantation and results stayed very similar (Figure S5A, B & C). This is not due to a proliferation rate change (Figure 8D). In comparison, such an immune rejection in immunocompetent BALB/c mice, which can be partially reversed by cadherin-11 overexpression, was not observed in immunodeficient NOD/SCID mice with the implantation of the same 4T1-luc cells (Figure 8E, F & G). Again, similarly to 4T1 cells without luciferase in BALB/c mice (Figure 7D-G), the tumor progression of 4T1-luc cells with cadherin-11 overexpression was much faster than 4T1-luc wildtype cells in NOD/SCID mice (Figure 8E, F & G). Such an interesting finding suggests that by hijacking mouse stromal fibroblasts in vivo by mouse cadherin-11-to-cadherin-11 adhesions, 4T1-luc cells with stable mouse cadherin-11 overexpression are able to significantly escape from the immune surveillance of the mouse immune system recognizing the firefly luciferase as a foreign antigen, which significantly prevents in vivo immune rejection against such cancer implants.

**Figure 8.**
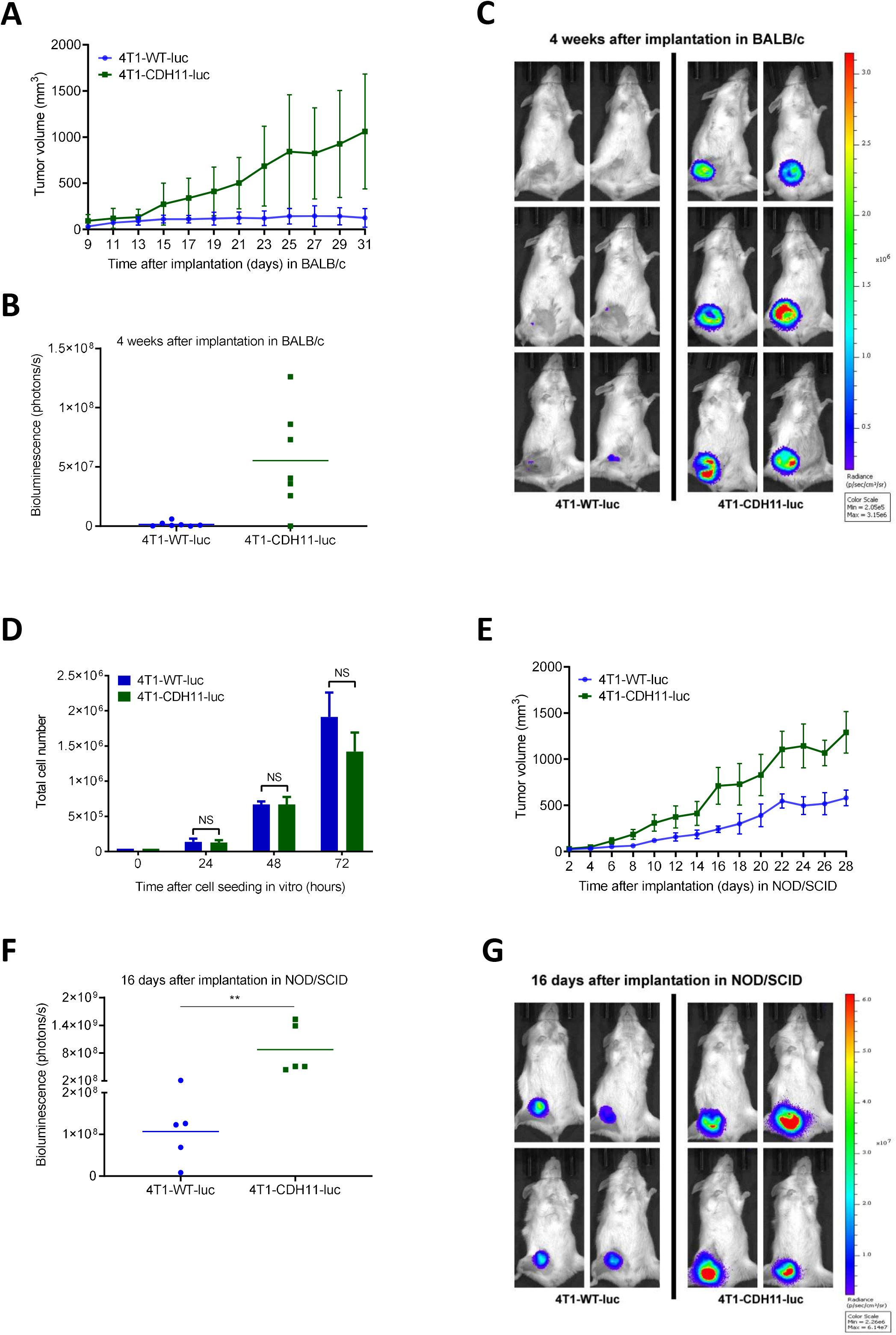
Strong immune rejection against tumor implants is detected in BALB/c mice implanted with 4T1-luc cells but is prevented by cadherin-11 overexpression in 4T1-luc cell implants. 1 × 10^6^ of 4T1 mouse triple negative breast cancer cells expressing firefly luciferase with or without CDH11 overexpression were implanted into the left fourth mammary fat pad of immunocompetent BALB/c mice (A-C) or NOD/SCID mice (E-G). A. Comparison of whole tumor growth volume between the 4T1-WT-luc cells implantation group and the 4T1-CDH11-luc cells implantation group in BALB/c mice. B. Comparison of cancer growth (as detected by the firefly luciferase bioluminescence) between the 4T1-WT-luc cells implantation group and the 4T1-CDH11-luc cells implantation group at 4 weeks after implantation in BALB/c mice. Data were presented as mean ± SD (n=7). C. Representative in vivo bioluminescence images from (B). D. Comparison of 4T1-WT-luc cell and 4T1-CDH11-luc cell proliferation in 2D culture in vitro. Same number of cells (46,000 cells) of each group were seeded in one well of a 6-well plate. Cell number was counted at 24 hrs, 48 hrs & 72 hrs (n=3 for each cell group at each time point) after cell seeding. NS, not significant. Note all cells were still not confluent at 72 hrs in each well of a 6-well plate. E. Comparison of whole tumor growth volume between the 4T1-WT-luc cells implantation group and the 4T1-CDH11-luc cells implantation group in NOD/SCID mice. F. Comparison of cancer growth (as detected by the firefly luciferase bioluminescence) between the 4T1-WT-luc cells implantation group and the 4T1-CDH11-luc cells implantation group at 4 weeks after implantation in NOD/SCID mice. Data were presented as mean ± SD (n=5). G. Representative *in vivo* bioluminescence images from (F).

## Discussion

Our findings firstly suggest that EMT is the origin but may not be the direct cause for cancer progression in vivo. Current prevailing EMT theory suggests epithelial cells undergo EMT so as to acquire the mesenchymal cell characteristics of being able to migrate, invade and metastasize on their own ^3^. For example, the two widely used human triple negative breast cancer cell lines, MDA-MB-231 and BT549, have been recognized as very migratory and invasive cancer cells ^53^. But these findings were previously observed mostly on 2D in vitro cell culture or Boyden Chamber based transwell cell migration and invasion assays. With the much-improved 3D ex vivo spheroid assays, we have observed that these cells are not very invasive on their own comparing to when they are cocultured with fibroblasts. Meanwhile, the previous in vivo findings of MDA-MB-231 and BT549 cells being invasive and metastatic in vivo were observed without considering their ideal physical interactions with fibroblasts in vivo by xenografting these human cancer cells expressing human cadherins into mouse stroma full of mouse cadherins ^44,45^. The cadherin biology studies have suggested that optimal cadherin-to-cadherin binding is achieved by “homophilic binding” between same types of cadherins not across species ^31,54^. Here we have determined that the “cadherin switch” from E-cadherin to cadherin-11 readily grants cancer cells the capability of hijacking fibroblasts, which significantly promotes cancer progression both in vitro and in vivo. Another example, the long-time deemed very aggressive 4T1 cells actually have only undergone partial EMT without cadherin switch. The only cadherin that 4T1 cells endogenously express is still E-cadherin but not any other mesenchymal cadherins (Figure S4A) ^51^, although they are already very migratory, invasive and aggressive both in vitro and in vivo ^49^. But, when we overexpressed cadherin-11 in 4T1 cells, they became even more aggressive and metastatic in vivo with significantly shorting mouse survival. Thus, we hereby provide significant updates to the EMT theory that the whole process of EMT may not significantly promote cancer progression itself, but the cadherin switch of the EMT process that enhances the interaction between cancer cells and the stroma, such a post-EMT event, might be the direct cause for enhanced cancer progression in human. Such cancer cell physical interactions with fibroblasts in such a short distance, the length of two extracellular regions of cadherin-11 proteins that are bound in trans ^55,56^, may play further roles in promoting cancer cell proliferation, differentiation, escape from immune surveillance (Figure 8), drug resistance and immunotherapy resistance beyond the studies that we have performed here.

Our study further suggests that EMT is sufficient and MET may not be required for cancer homing. Recent studies have observed the plasticity of EMT in vivo and suggest that the reverse process MET promotes cancer homing and outgrowth at metastatic sites ^57^. Stable knockdown of PRRX1 or TWIST or both, two of the key EMT-TFs, induced cancer colonization and outgrowth in lung ^7,8^. These studies have suggested that a partial or even complete reverse of EMT by the MET process might be involved in cancer homing and outgrowth in metastatic sites and have raised controversies and debates on the EMT theory ^3,6^. In comparison, we have observed that simply overexpressing the mesenchymal cadherin-11 in 4T1 cells, who endogenously only express E-cadherin, to enhance their adhesions to stromal fibroblasts is sufficient to promote their growth in situ, progression, metastasis and outgrowth in metastatic sites, such as lymph nodes (Figure 7). Our study suggests that the cadherin-11 mediated cancer cell hijacking fibroblasts is enough to promote cancer homing and MET is not required in such cases.

EMT cancer cells may be way smarter than we ever thought. For the cell invasion research field, in addition to the current knowledge of cancer cell self-invasion, cancer cell following fibroblast-made matrix tunnel invasion ^17,18^, cancer cell coupling with macrophage invasion by an EGF/CSF1 paracrine signaling loop ^58–60^, we have discovered a novel cancer cell invasion mechanism of cadherin-11 homophilic adhesion mediated cancer cell hijacking fibroblast invasion. Such cell “hijacking” is achieved by EMT cancer cells, especially triple negative breast cancer cells, expressing fibroblast specific protein cadherin-11 ^26,32^. This finding might help untangle the puzzle of why cadherin-11 expressing cancer cells are always more metastatic in vivo ^27,44^. Further, our study reveals such an alternative invasion mode when cadherin-11 positive cancer cells are surrounded by fibroblasts. Instead of being trapped and constrained by these neighboring giant cells, cancer cells hijack them by cadherin-11 and go with them. Interestingly, EMT cancer cells can do sole invasion by themselves, but when they manage to adhere to fibroblasts by engaging the cadherin-11-to-cadherin-11 adhesions, they gave up their own mesenchymal invasion but rather take the fibroblast “train” for a co-invasion, which may significantly save tremendous energy that cancer cells have to spend in doing mesenchymal invasion on their own. Such a smart energy saving strategy may further let cancer cells spend their precious energy in other valuable activities such as homing, proliferation and inducing angiogenesis.

Lastly, previous studies have focused on the extracellular intercell signalings between the cancer cells and the stromal fibroblasts in the “Seed and Soil” theory of cancer metastasis and homing ^10^. Our study suggests that cadherin-11 can be the key molecule that readily engages EMT cancer cells to the surrounding stroma for enhanced survival, local invasion, distal metastasis, escape from immune surveillance and much more.

## MATERIALS AND METHODS

### Cells and proteins

Primary human fibroblasts were isolated from synovial tissue ^23,32^ or purchased from ATCC (Manassas, VA). These cells were used for experiments between passages 5 to 10. MDA-MB-231, BT549, 4T1, NIH3T3, 293T cells were all purchased from ATCC (Manassas, VA). All cells were cultured in DMEM, supplemented with 10% fetal bovine serum (FBS), L-glutamine, and penicillin/streptomycin. Fibroblast growth factor (FGF) was added for maintaining primary human fibroblasts. All cell culture reagents were obtained from Thermo Fisher Scientific (Carlsbad, CA).

### Animal studies

All experiments in mice were performed in accordance with relevant institutional guidelines and regulations and were subject to a protocol approved by Department of Health, Hong Kong. NOD/SCID female mice and BALB/c female mice were purchased from the Laboratory Animal Services Centre, the Chinese University of Hong Kong. Animals were housed 4-6 per cage under a light-dark (12 h/12 h) cycle with ad libitum access to water and food. Mice were 6-8 weeks old at time of tumor implantation. Cells at the exponential phase of growth were harvested and suspended with a matrigel (1.5 mg/ml, Cultrex® BME, Trevigen, Gaithersburg, MD) and collagen (0.35 mg/ml, Corning® Collagen I, Corning, NY) mixture. Mice were injected with 200 μl of cell suspension containing 1×10^6^ or 2×10^6^ cells into the left fourth abdominal fat pad by injection at the base of the nipple ^61^. During injection mice were kept under inhalation anesthesia induced by 3% (V/V) and maintained at 0.8 - 1% (V/V) mixture of isoflurane/oxygen through facial masks. Bioluminescence of tumors was measured every week on an IVIS Lumina XR imaging system (Perkin Elmer, Waltham, MA) after intraperitoneal injection of D-luciferin (150 mg/kg, 200 μl). Images were capture every minute for continuous 15 mins and analyzed by the Living Image® software (Perkin Elmer, Waltham, MA, USA). Tumor growth was monitored by measuring tumor length (L) and width (W) using Vernier calipers weekly (NOD/SCID mice) or every 2-3 days (BALB/c mice). Tumor volume (V) was calculated as [V = (L × W^2^)/2].

### PDX establishment

The human breast cancer tissues were obtained from patients with informed consent who had surgery within 3 hours at the Department of Breast Surgery, Fudan University Shanghai Cancer Center (Shanghai, China). Tumor fragments of 30 to 60 mm^3^ in volume were subcutaneously implanted in 6-week old NOG mice (CIEA, Japan) weighing from 18 to 25 grams.

### Micro-MRI (Magnetic Resonance Imaging)

All images were acquired using a 3 Tesla integrated small animal PET/MR scanner (nanoScan, Mediso Medical Imaging Systems Ltd, Budapest, Hungary). Mice were anesthetized via inhalation of 5 % isoflurane/oxygen gas mixture and placed on a pre-warmed mouse bed in the prone position. Mice were maintained using 2% isoflurane/oxygen during the imaging acquisition periods and were tightly wrapped together with the bed using a paper tape to reduce respiratory motion artifacts. For tumor detection, scans were performed in the coronal plane using a respiratory triggered T2-weighted 2D fast spin echo sequence with the following parameters: TR/TE 4000/71.4 ms; slice thickness 0.9 mm; in-plane matrix 336 × 224; in-plane resolution 0.27 × 0.27 mm2; five signal acquisition. The average acquisition time was approximately 12 min, depending on the respiration rate of the mice.

### Plasmid, siRNA, and transfection

Mouse cadherin-11 expression plasmid was from Sino biological Inc. (Wayne, PA). pGL4.51[luc2/CMV/Neo] vector encoding the firefly luciferase reporter gene luc2 was from Promega (Madison, WI). All siRNAs and shRNAs were from Thermo Fisher Scientific (Carlsbad, CA) or Dharmacon (Lafayette, CO, USA). Lipofectamine™ LTX, Lipofectamine™ 2000, and Lipofectamine™ 3000 reagents (Invitrogen, Carlsbad, CA) were used for transfecting expression plasmids. Lipofectamine™ RNAiMAX reagent (Invitrogen, Carlsbad, CA) was used for siRNA transfection. Lentiviruses carrying shRNA with GFP reporter were generated in 293T cells by the Trans-Lentiviral shRNA Packaging System (Dharmacon, Lafayette, CO). Lentiviruses were then collected and transferred to infect shRNA targeting cells. Stable shRNA silencing cell clones were selected by relevant antibiotics followed by cell sorting by the FACSAria III cell sorter flow cytometer (BD Biosciences, San Jose, CA).

### Quantitative real-time polymerase chain reaction

Total RNAs were extracted by the Aurum ™ Total RNA mini kit (Bio-Rad Laboratories Inc., Hercules, CA, USA) and subject to qRT-PCR assays by the Power SYBR® Green RNA-to-Ct™ 1-Step Kit (Thermo Fisher Scientific, Waltham, MA) to measure molecule expression levels. Expression level of glyceraldehyde-3-phosphate dehydrogenase (GAPDH) was used as reference control. qRT-PCR primers were from Sino Biological (Wayne, PA) or Genetimes ExCell (Hong Kong). All qRT-PCR experiments were run on the ViiA™ 7 Real-Time PCR System (Life Technologies, Thermo Fisher Scientific, Waltham, MA)

### Antibodies

Mouse anti-cadherin-11 (clone 3H10 and 23C6) antibody has been described previously ^26^. Mouse anti-cadherin-11 (clone 16A) antibody was from Millipore (Billerica, MA). Mouse anti-cadherin-11 (clone 5B2H5), Alexa Fluor conjugated anti-mouse, anti-rat and anti-rabbit secondary antibodies and Alexa Fluor conjugated phalloidin were obtained from Invitrogen (Carlsbad, CA).

### Microscopy

For immunofluorescence microscopy, cells were labeled with a primary antibody for 1 hour at room temperature or 16 hours at 4°C, followed by secondary antibody labeling for 1 hour, and then mounted on slides with FluorSave™ Reagent (Calbiochem, San Diego, CA). For confocal microscopy, images were captured on various confocal microscopes at the Brigham & Women’s Hospital Confocal Microscopy Core Facility, the Analytical Imaging Facility of the Albert Einstein College of Medicine, the Nikon Imaging Center at Harvard Medical School, the Integrated Imaging Center at Johns Hopkins University, the Multiphoton Microscopy Core Facility at The MD Anderson Cancer Center and the Confocal Microscopy Core Facility at Hong Kong Baptist University. Confocal microscopy was performed by scanning multiple confocal layers along the z-axis. For live-cell imaging, cells were grown in 35 mm glass-bottomed Fluorodishes (World Precision Instruments, Sarasota, Florida), and then mounted onto the confocal microscope equipped with a heating chamber warmed to 37 °C and supplied with 10% CO_2_. Sections of confocal images were scanned along the cell z axis every 1 min. After imaging, 3-D stacks of images at each time point were projected into a 2D image. Image reconstruction and quantitative colocalization analyses were performed using MetaMorph imaging software version 7.6.1.0 (Universal Imaging, Chesterfield, PA), Leica LAS X software (Leica Microsystems GmbH, Wetzlar, Germany), Zeiss Zen microscope software (Carl Zeiss Microscopy GmbH, Jena, Germany), ImageJ software or Fiji software, etc.

### 3-D spheroid cell invasion assay and microfluidic devices

Cells were pre-labeled by using the Vybrant™ Multicolor Cell-Labeling Kit (DiO, DiI, DiD Solutions) (Invitrogen, Carlsbad, CA). Each spheroid was made by resuspending 8,000 cells in 15 μl media with 0.24% methyl cellulose (4,000 cpi, Sigma-Aldrich, St. Louis, MO) by the hanging drop method ^34^. Each spheroid was then embedded in 400 μl extracellular matrix (ECM) with 6 mg/ml matrigel (Cultrex® BME, Trevigen, Gaithersburg, MD) and 1.4 mg/ml collagen (Corning® Collagen I, Corning, NY). After the ECM was solidified, 200 μl liquid media were placed on top to cover the ECM. The final serum concentration in all spheroid assay experiments was 6.7% except for the cytokine stimulation spheroid experiments, in which the final serum concentration was 1.3%. The 3-D spheroid made by this hanging drop method is like a 3-D pancake shape with long X-Y axes but a shorter Z axis, which is perfect for confocal imaging. Cell invasion was quantitated by the ratio of the protrusion area over the core area (Figure 2A). The microfluidic devices were manufactured by following the method that was described previously ^62^. Further details were described in Figure 2D.

### Western blotting

Cells were lysed in buffer (1% Triton X-100, 50 mM HEPES pH 7.4, 150 mM NaCl, 5 mM EDTA, 5 mM EGTA, 20 mM NaF, 20 mM sodium pyrophosphate, 1 mM PMSF, 1 mM Na3VO4). Samples were then separated by SDS-PAGE gels, transferred to PVDF membrane, and visualized by chemiluminescence (Thermo Fisher Scientific, Rockford, IL).

### Statistics

Data are presented as means ± SD (standard deviation). Statistical analysis was performed using the two-tailed Student’s t test with GraphPad Prism 7, version 7.01 (GraphPad Software, Inc., San Diego, CA). P values less than 0.05 were considered as statistically significant, while those less than 0.01 were considered as highly significant (marked by stars in figures).

## Acknowledgements

This work was supported by Hong Kong University Grants Committee/ Research Grants Council Grants ECS22103319 and GRF17210618 to Z.G.; Hong Kong Baptist University Grants Startup/38-402-94, FRG2/17-18/039 and SGT2/18-19/018 to Z.G.; The National Institute of Health Grants P01CA100324 and R21NS087624 to J.E.S.; The National Institute of Health Grant P30CA013330 to the Analytical Imaging Facility of the Albert Einstein College of Medicine; Hong Kong University Grants Committee/Research Grant Council Grant CRFC7018-14E to The MicroPET-MRI Laboratory of The University of Hong Kong.

We thank The Brigham & Women’s Hospital Confocal Microscopy Core Facility, The Analytical Imaging Facility of the Albert Einstein College of Medicine, The Nikon Imaging Center at Harvard Medical School, The Integrated Imaging Center at Johns Hopkins University, The Multiphoton Microscopy Core Facility at The MD Anderson Cancer Center and The MicroPET-MRI Laboratory of The University of Hong Kong for their generous helps and supports in imaging.

## Author contributions

Z.G. raised the hypothesis, conceptualized and designed the project, performed some of the experiments. W.K., Y.F. and Y.D. contributed equally to performing experiments. W.K. and E.A.T. contributed to writing the manuscript. Y.H.H. and M.C.H. discovered the correlation between cadherin-11 expression and triple negative breast cancers. E.A.T., H.Y., J.L., K.Y., J.Y. helped with experiments. K.V.T. and P.L.K. provided collaboration in Micro-MRI experiments. S.A. and P.F. provided collaboration in imaging method designing. B.S.W. and K.K. provided collaboration in microfluidics experiments. X.H., T.L., Y.G. and J.Z. provided collaboration in PDX experiments. J.J.Z., A.L. and M.B.B. provided collaborations and supports. J.E.S. provided collaborations and supports, helped with designing experiments and contributed to writing the manuscript. Z.G. supervised all aspects of the project and wrote the manuscript.

## References

1. Brabletz, T. To differentiate or not--routes towards metastasis. Nat Rev Cancer 12, 425–436 (2012).

2. Thiery, J.P., Acloque, H., Huang, R.Y. & Nieto, M.A. Epithelial-mesenchymal transitions in development and disease. Cell 139, 871–890 (2009).

3. Brabletz, T., Kalluri, R., Nieto, M.A. & Weinberg, R.A. EMT in cancer. Nat Rev Cancer 18, 128–134 (2018).

4. Thiery, J.P. & Sleeman, J.P. Complex networks orchestrate epithelial-mesenchymal transitions. Nature reviews. Molecular cell biology 7, 131–142 (2006).

5. Kalluri, R. & Weinberg, R.A. The basics of epithelial-mesenchymal transition. J Clin Invest 119, 1420–1428 (2009).

6. Nieto, M.A., Huang, R.Y., Jackson, R.A. & Thiery, J.P. Emt: 2016. Cell 166, 21–45 (2016).

7. Ocana, O.H., et al. Metastatic colonization requires the repression of the epithelial-mesenchymal transition inducer Prrx1. Cancer Cell 22, 709–724 (2012).

8. Tsai, J.H., Donaher, J.L., Murphy, D.A., Chau, S. & Yang, J. Spatiotemporal regulation of epithelialmesenchymal transition is essential for squamous cell carcinoma metastasis. Cancer Cell 22, 725–736 (2012).

9. Li, G., et al. Function and regulation of melanoma-stromal fibroblast interactions: when seeds meet soil. Oncogene 22, 3162–3171 (2003).

10. Mueller, M.M. & Fusenig, N.E. Friends or foes - bipolar effects of the tumour stroma in cancer. Nat Rev Cancer 4, 839–849 (2004).

11. Turley, S.J., Cremasco, V. & Astarita, J.L. Immunological hallmarks of stromal cells in the tumour microenvironment. Nat Rev Immunol 15, 669–682 (2015).

12. Pallangyo, C.K., Ziegler, P.K. & Greten, F.R. IKKbeta acts as a tumor suppressor in cancer-associated fibroblasts during intestinal tumorigenesis. The Journal of experimental medicine 212, 2253–2266 (2015).

13. Koliaraki, V., Pasparakis, M. & Kollias, G. IKKbeta in intestinal mesenchymal cells promotes initiation of colitis-associated cancer. The Journal of experimental medicine 212, 2235–2251 (2015).

14. Karnoub, A.E., et al. Mesenchymal stem cells within tumour stroma promote breast cancer metastasis. Nature 449, 557–563 (2007).

15. Perentes, J.Y., et al. In vivo imaging of extracellular matrix remodeling by tumor-associated fibroblasts. Nat Methods 6, 143–145 (2009).

16. Duda, D.G., et al. Malignant cells facilitate lung metastasis by bringing their own soil. Proc Natl Acad Sci U S A 107, 21677–21682 (2010).

17. Gaggioli, C., et al. Fibroblast-led collective invasion of carcinoma cells with differing roles for RhoGTPases in leading and following cells. Nature cell biology 9, 1392–1400 (2007).

18. Scott, R.W., et al. LIM kinases are required for invasive path generation by tumor and tumorassociated stromal cells. The Journal of cell biology 191, 169–185 (2010).

19. Del Pozo Martin, Y., et al. Mesenchymal Cancer Cell-Stroma Crosstalk Promotes Niche Activation, Epithelial Reversion, and Metastatic Colonization. Cell Rep 13, 2456–2469 (2015).

20. Avgustinova, A., et al. Tumour cell-derived Wnt7a recruits and activates fibroblasts to promote tumour aggressiveness. Nat Commun 7, 10305 (2016).

21. Otomo, R., et al. TSPAN12 is a critical factor for cancer-fibroblast cell contact-mediated cancer invasion. Proc Natl Acad Sci U S A 111, 18691–18696 (2014).

22. Okazaki, M., et al. Molecular cloning and characterization of OB-cadherin, a new member of cadherin family expressed in osteoblasts. J Biol Chem 269, 12092–12098 (1994).

23. Chang, S.K., et al. Cadherin-11 regulates fibroblast inflammation. Proc Natl Acad Sci U S A 108, 8402–8407 (2011).

24. Roosen, A., et al. Cadherin-11 up-regulation in overactive bladder suburothelial myofibroblasts. J Urol 182, 190–195 (2009).

25. Shibata, T., Ochiai, A., Gotoh, M., Machinami, R. & Hirohashi, S. Simultaneous expression of cadherin-11 in signet-ring cell carcinoma and stromal cells of diffuse-type gastric cancer. Cancer Lett 99, 147–153 (1996).

26. Valencia, X., et al. Cadherin-11 provides specific cellular adhesion between fibroblast-like synoviocytes. The Journal of experimental medicine 200, 1673–1679 (2004).

27. Pishvaian, M.J., et al. Cadherin-11 is expressed in invasive breast cancer cell lines. Cancer research 59, 947–952 (1999).

28. Tomita, K., et al. Cadherin switching in human prostate cancer progression. Cancer research 60, 3650–3654 (2000).

29. Sarrio, D., et al. Epithelial-mesenchymal transition in breast cancer relates to the basal-like phenotype. Cancer research 68, 989–997 (2008).

30. Takeichi, M. The cadherins: cell-cell adhesion molecules controlling animal morphogenesis. Development 102, 639–655 (1988).

31. Yap, A.S., Gomez, G.A. & Parton, R.G. Adherens Junctions Revisualized: Organizing Cadherins as Nanoassemblies. Dev Cell 35, 12–20 (2015).

32. Lee, D.M., et al. Cadherin-11 in synovial lining formation and pathology in arthritis. Science 315, 1006–1010 (2007).

33. Gu, Z. 0.1 kilopascal difference for mechanophenotyping: soft matrix precisely regulates cellular architecture for invasion. Bioarchitecture 4, 116–118 (2014).

34. Veelken, C., Bakker, G.J., Drell, D. & Friedl, P. Single cell-based automated quantification of therapy responses of invasive cancer spheroids in organotypic 3D culture. Methods 128, 139–149 (2017).

35. Hung, W.C., et al. Distinct signaling mechanisms regulate migration in unconfined versus confined spaces. The Journal of cell biology 202, 807–824 (2013).

36. Gu, Z., et al. Soft matrix is a natural stimulator for cellular invasiveness. Molecular biology of the cell 25, 457–469 (2014).

37. Ramis-Conde, I., Drasdo, D., Anderson, A.R. & Chaplain, M.A. Modeling the influence of the E-cadherin-beta-catenin pathway in cancer cell invasion: a multiscale approach. Biophys J 95, 155–165 (2008).

38. Smutny, M., et al. Friction forces position the neural anlage. Nature cell biology 19, 306–317 (2017).

39. Kiener, H.P., et al. Cadherin 11 promotes invasive behavior of fibroblast-like synoviocytes. Arthritis Rheum 60, 1305–1310 (2009).

40. Barretina, J., et al. The Cancer Cell Line Encyclopedia enables predictive modelling of anticancer drug sensitivity. Nature 483, 603–607 (2012).

41. Pharmacogenomic agreement between two cancer cell line data sets. Nature 528, 84–87 (2015).

42. Uhlen, M., et al. A pathology atlas of the human cancer transcriptome. Science 357(2017).

43. Nagy, A., Lanczky, A., Menyhart, O. & Gyorffy, B. Validation of miRNA prognostic power in hepatocellular carcinoma using expression data of independent datasets. Sci Rep 8, 9227 (2018).

44. Assefnia, S., et al. Cadherin-11 in poor prognosis malignancies and rheumatoid arthritis: common target, common therapies. Oncotarget 5, 1458–1474 (2014).

45. Satriyo, P.B., et al. Cadherin 11 Inhibition Downregulates beta-catenin, Deactivates the Canonical WNT Signalling Pathway and Suppresses the Cancer Stem Cell-Like Phenotype of Triple Negative Breast Cancer. J Clin Med 8(2019).

46. Kojima, M., et al. Human subperitoneal fibroblast and cancer cell interaction creates microenvironment that enhances tumor progression and metastasis. PloS one 9, e88018 (2014).

47. Murata, T., Mekada, E. & Hoffman, R.M. Reconstitution of a metastatic-resistant tumor microenvironment with cancer-associated fibroblasts enables metastasis. Cell Cycle 16, 533–535 (2017).

48. Chatterjee, S., et al. Breast Cancers Activate Stromal Fibroblast-Induced Suppression of Progenitors in Adjacent Normal Tissue. Stem Cell Reports 10, 196–211 (2018).

49. Pulaski, B.A. & Ostrand-Rosenberg, S. Mouse 4T1 breast tumor model. Curr Protoc Immunol Chapter 20, Unit 20 22 (2001).

50. Bouquet, F., et al. TGFbeta1 inhibition increases the radiosensitivity of breast cancer cells in vitro and promotes tumor control by radiation in vivo. Clin Cancer Res 17, 6754–6765 (2011).

51. Wagenblast, E., et al. A model of breast cancer heterogeneity reveals vascular mimicry as a driver of metastasis. Nature 520, 358–362 (2015).

52. Baklaushev, V.P., et al. Luciferase Expression Allows Bioluminescence Imaging But Imposes Limitations on the Orthotopic Mouse (4T1) Model of Breast Cancer. Sci Rep 7, 7715 (2017).

53. Cui, B., et al. Targeting ROR1 inhibits epithelial-mesenchymal transition and metastasis. Cancer research 73, 3649–3660 (2013).

54. Shapiro, L. & Weis, W.I. Structure and biochemistry of cadherins and catenins. Cold Spring Harbor perspectives in biology 1, a003053 (2009).

55. Patel, S.D., et al. Type II cadherin ectodomain structures: implications for classical cadherin specificity. Cell 124, 1255–1268 (2006).

56. Hong, S., Troyanovsky, R.B. & Troyanovsky, S.M. Cadherin exits the junction by switching its adhesive bond. The Journal of cell biology 192, 1073–1083 (2011).

57. Alderton, G.K. Metastasis: Epithelial to mesenchymal and back again. Nat Rev Cancer 13, 3 (2013).

58. Condeelis, J. & Pollard, J.W. Macrophages: obligate partners for tumor cell migration, invasion, and metastasis. Cell 124, 263–266 (2006).

59. Wyckoff, J., et al. A paracrine loop between tumor cells and macrophages is required for tumor cell migration in mammary tumors. Cancer research 64, 7022–7029 (2004).

60. Harney, A.S., et al. Real-Time Imaging Reveals Local, Transient Vascular Permeability, and Tumor Cell Intravasation Stimulated by TIE2hi Macrophage-Derived VEGFA. Cancer Discov 5, 932–943 (2015).

61. Iorns, E., et al. A new mouse model for the study of human breast cancer metastasis. PloS one 7, e47995 (2012).

62. Hung, W.C., et al. Confinement Sensing and Signal Optimization via Piezo1/PKA and Myosin II Pathways. Cell Rep 15, 1430–1441 (2016).

