## Supplementary Material for "Post-EMT: Cadherin-11 mediates cancer hijacking fibroblasts"

**SUPPLEMENTAL FIGURE AND VIDEO LEGEND**

**Figure S1. Cadherin-11 expressing cancer cells hijacking fibroblasts in a large population.**
A. The schematic to describe the large-population cell invasion assay by multiphoton imaging. B. Primary human fibroblasts were pre-labeled by DiI (red) and MDA-MB-231 cells were pre-labeled by DiO (green). Cells were mixed by the 1:1 ratio in cell numbers and were subjected to the cell invasion assay in (A). A 200 µm thick Z-stack were scanned by the multiphoton microscope from the top aiming at the invasion leading cells and a 3-D image was reconstructed by the Zeiss Zen software.

**Figure S2. Cadherin-11 downregulation.**
A & B. MDA-MB-231 cells transfected with negative control siRNA or the siRNA against cadherin-11 were cultured for 48 hours. Cells were then lysed and whole cell lysates were subjected to SDS-PAGE and western blotting. Cadherin-11 proteins were blotted and quantified (B). *, $P < 0.05$. C. The table explaining the effects of cadherin-11 downregulation in Figure 3A.

**Figure S3. The cancer cell hijacking fibroblast invasion is cadherin-11 dependent.**
A. MDA-MB-231 cells and fibroblasts coculture (1:1) spheroid with negative control siRNA or CDH11 siRNA were subjected to the 3D spheroid cell invasion assay as in Figure 3C. Total cell numbers maintained the same in every spheroid. Confocal Z-stacks were scanned from the top of the spheroid to the bottom. All 2-D images shown were from 3-D Z-stacks maximum projections. B, C & D. Quantitation for images as in (A). Experiments were repeated 6 times (n=6). *, $P < 0.05$.

**Figure S4. Gene expression of Cdh1 to Cdh11 in 4T1 cells.** A. The gene expression levels of mouse cadherin family members were analyzed based on the normalized RNAseq data (dataset ID: GSE63180) (Wagenblast et al., 2015) from the NCBI GEO (Gene Expression Omnibus, National Center for Biotechnology Information, U.S. National Library of Medicine) database. Data were presented as mean ± SD (n=46, duplicates of 23 single clonal 4T1 cell lines established by single cell FACS sorting), representing gene expression folds over Cdh2 average expression level (normalized to 1).

**Figure S5. A repetitive experiment as in Figure 8 using completely different batch of stable CDH11 overexpression cells.** $1 \times 10^6$ of 4T1 mouse triple negative breast cancer cells expressing firefly luciferase with or without CDH11 overexpression were implanted into the left fourth mammary fat pad of immunocompetent BALB/c mice (A-C). A. Comparison of whole tumor growth volume between the 4T1-WT-luc cells implantation group and the 4T1-CDH11-luc cells implantation group in BALB/c mice. B. Comparison of cancer growth (as detected by the firefly luciferase bioluminescence) between the 4T1-WT-luc cells implantation group and the 4T1-CDH11-luc cells implantation group at 4 weeks after implantation in BALB/c mice. Data were presented as mean ± SD (n=10). C. Representative *in vivo* bioluminescence images from (B).

**Video 1, 2 & 3.** Fibroblasts and MDA-MB-231 cells were pre-labeled as before. Fibroblasts alone (red, Video 1), MDA-MB-231 alone (blue, Video 2) or Fibroblasts and MDA-MB-231 (Video 3) together in a 1:1 ratio were subjected to the cell invasion assay as described in Figure 1C. The green fluorescence from the matrigel was omitted to clearly visualize the cells.

**Video 4.** Time-lapse zoom-in video from Video 3.

**Video 5.** Primary human fibroblasts were pre-labeled by DiI (red) and MDA-MB-231 cells were pre-labeled by DiO (green). Cells were subjected to the microfluidic invasion assay as described in Figure 2D to observe the cancer cell hijacking fibroblast co-invasion at single cell level.

**Fig. S1**

**A**

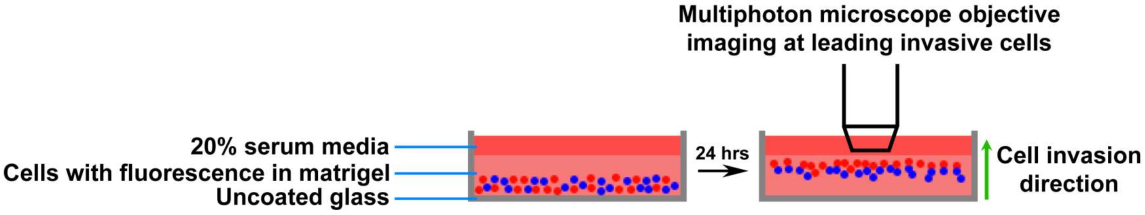
Multiphoton microscope objective
imaging at leading invasive cells
20% serum media
Cells with fluorescence in matrigel
Uncoated glass
24 hrs
Cell invasion
direction

**B**

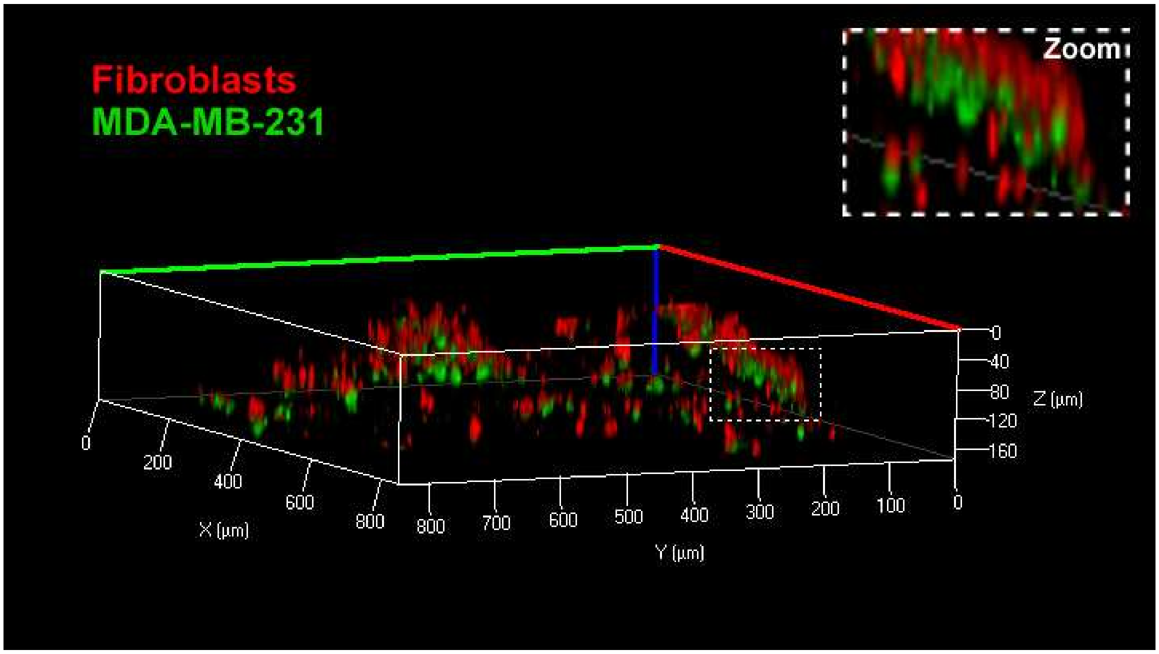
Fibroblasts
MDA-MB-231
Zoom
0
40
80
120
160
Z (µm)
0
200
400
600
800
X (µm)
800
700
600
500
400
300
200
100
0
Y (µm)

**Fig. S2**

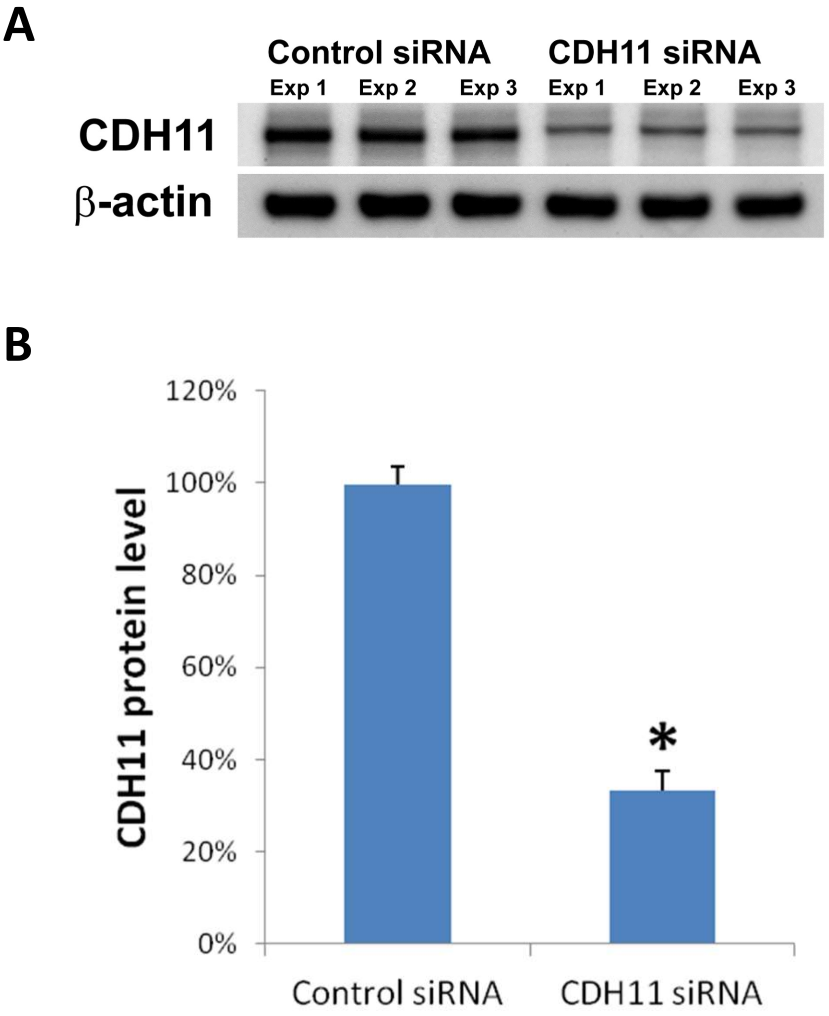


**C**

| | CDH11 friction reduction induced cell dissemination [A] | | CDH11 promoted cell invasion by an unknown mechanism [B] | | Overall cell invasion [A+B] | |
|---|---|---|---|---|---|---|
| | Control | CDH11 knockdown | Control | CDH11 knockdown | Control | CDH11 knockdown |
| Cancer cells | – | ↑ | – | ↑↓? | – | ↑ |
| Fibroblasts | – | ↑ | – | ↓↓↓ | – | ↓↓ |

**Fig. S3**

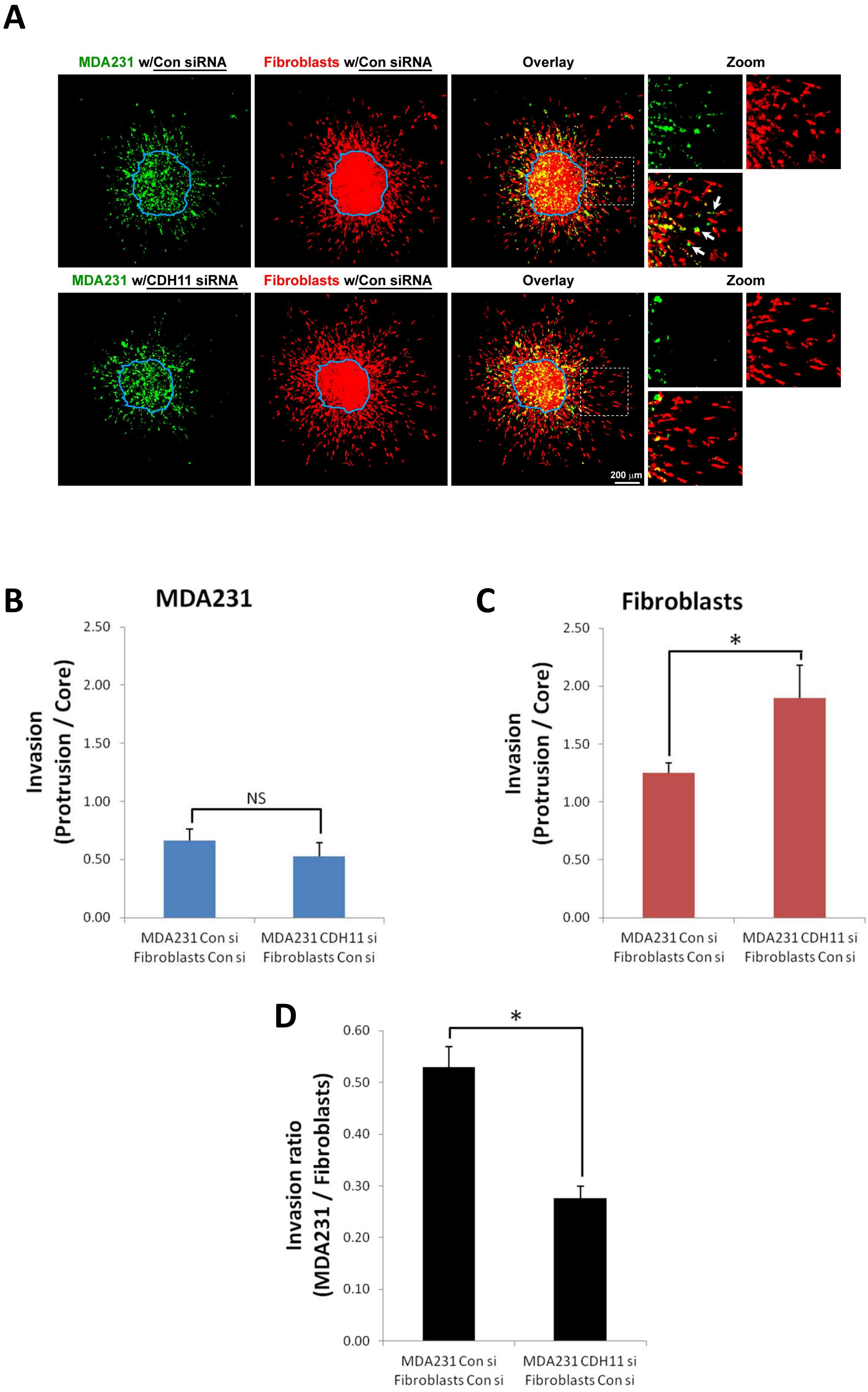

A
MDA231 w/Con siRNA
Fibroblasts w/Con siRNA
Overlay
Zoom
MDA231 w/CDH11 siRNA
Fibroblasts w/Con siRNA
Overlay
Zoom
200 μm
B
MDA231
Invasion (Protrusion / Core)
2.50
2.00
1.50
1.00
0.50
0.00
NS
MDA231 Con si Fibroblasts Con si
MDA231 CDH11 si Fibroblasts Con si
C
Fibroblasts
Invasion (Protrusion / Core)
2.50
2.00
1.50
1.00
0.50
0.00
*
MDA231 Con si Fibroblasts Con si
MDA231 CDH11 si Fibroblasts Con si
D
Invasion ratio (MDA231 / Fibroblasts)
0.60
0.50
0.40
0.30
0.20
0.10
0.00
*
MDA231 Con si Fibroblasts Con si
MDA231 CDH11 si Fibroblasts Con si

Fig. S4

A

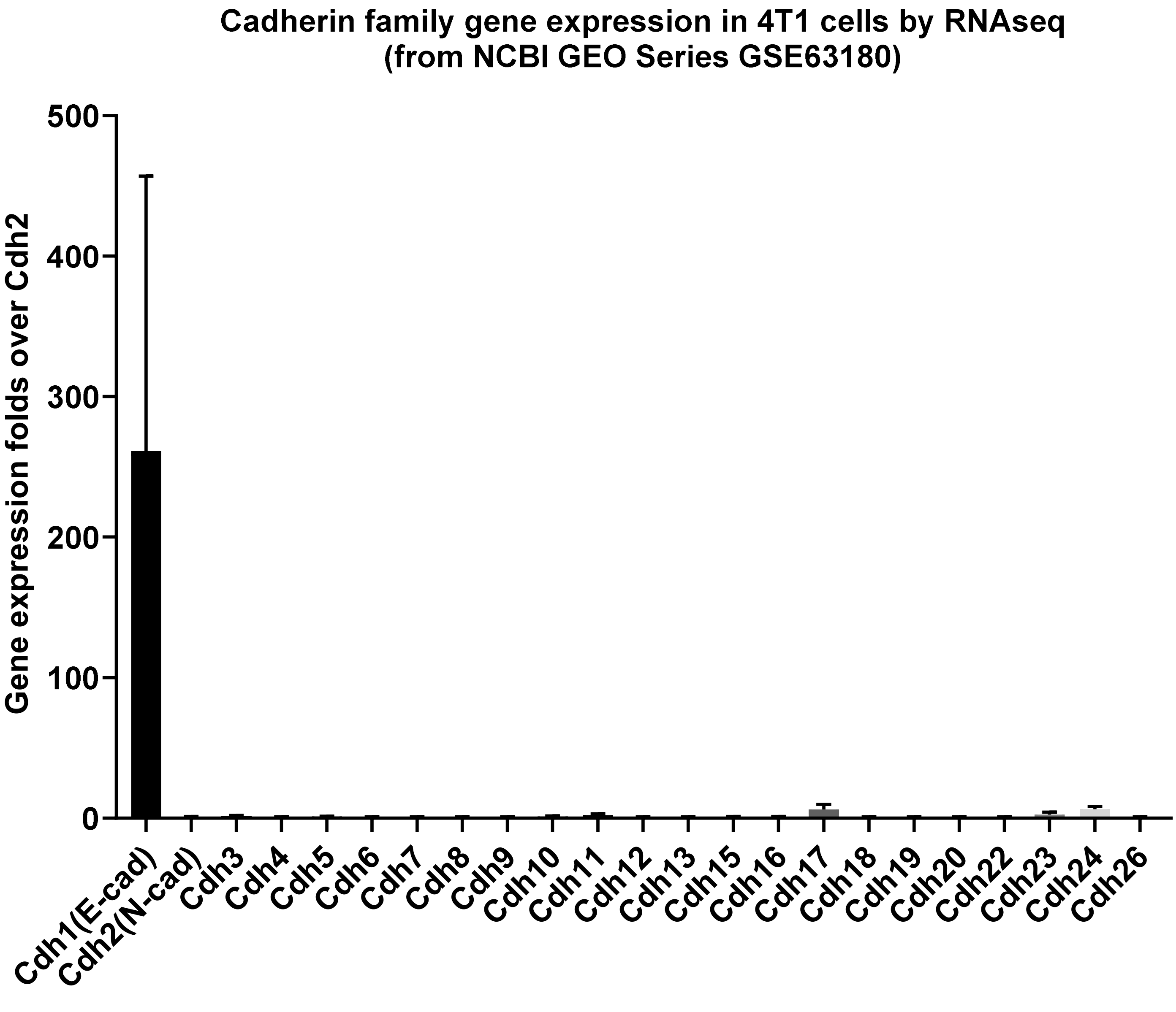

Cadherin family gene expression in 4T1 cells by RNAseq
(from NCBI GEO Series GSE63180)
Gene expression folds over Cdh2
500
400
300
200
100
0
Cdh1(E-cad)
Cdh2(N-cad)
Cdh3
Cdh4
Cdh5
Cdh6
Cdh7
Cdh8
Cdh9
Cdh10
Cdh11
Cdh12
Cdh13
Cdh15
Cdh16
Cdh17
Cdh18
Cdh19
Cdh20
Cdh22
Cdh23
Cdh24
Cdh26

Fig. S5

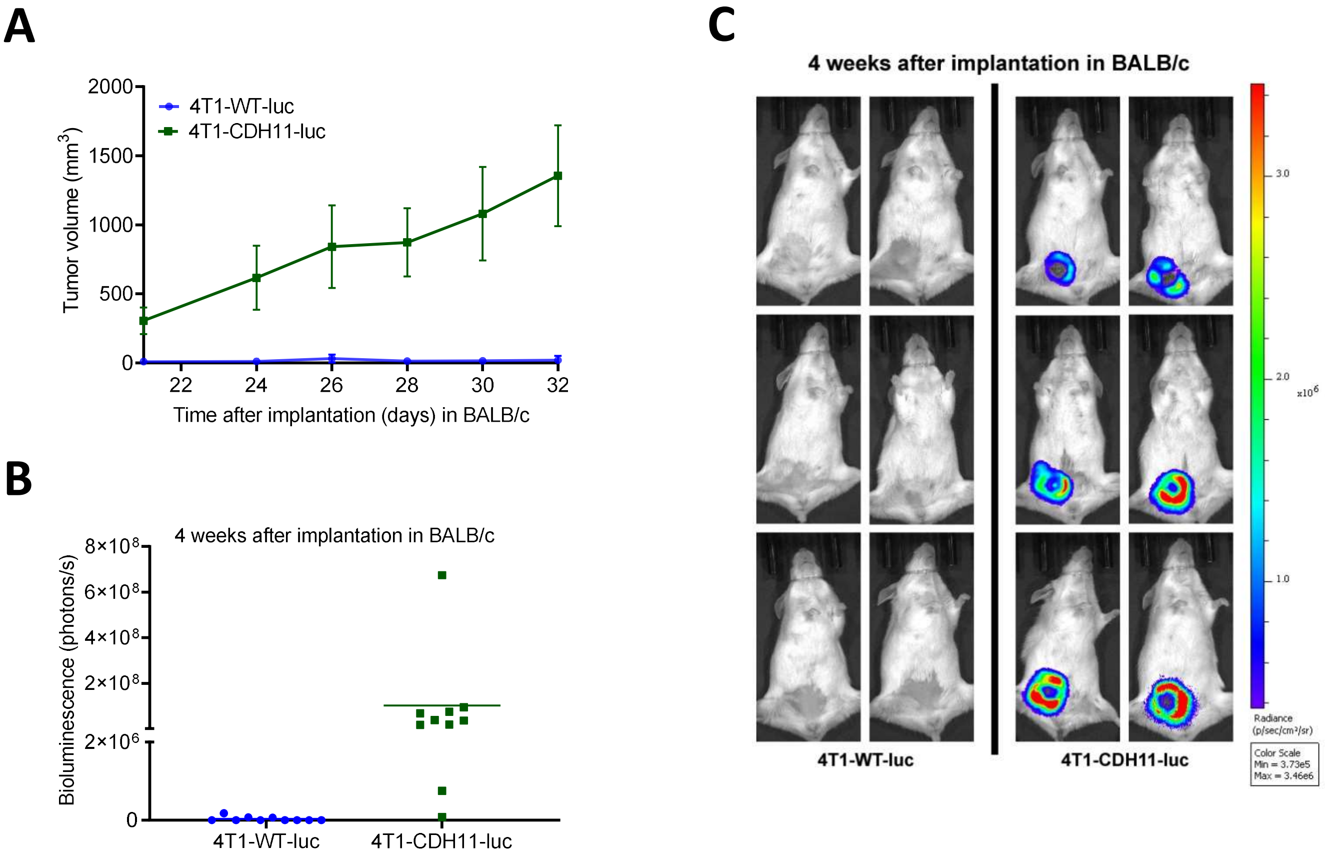

A
2000
1500
1000
500
0
Tumor volume (mm3)
4T1-WT-luc
4T1-CDH11-luc
22
24
26
28
30
32
Time after implantation (days) in BALB/c
B
4 weeks after implantation in BALB/c
8×10^8
6×10^8
4×10^8
2×10^8
2×10^6
0
Bioluminescence (photons/s)
4T1-WT-luc
4T1-CDH11-luc
C
4 weeks after implantation in BALB/c
3.0
2.0
x10^6
1.0
Radiance (p/sec/cm²/sr)
Color Scale Min = 3.73e5 Max = 3.46e6
4T1-WT-luc
4T1-CDH11-luc
